# RBPscan: A Quantitative, In Vivo Tool for Profiling RNA-Binding Protein Interactions

**DOI:** 10.1101/2025.01.03.631239

**Authors:** Dmitry A. Kretov, Owen Sanborn, Thora McIssac, Elaine Park, Imrat, Samuel Wu, Daniel Cifuentes

## Abstract

RNA-binding proteins (RBPs) are essential regulators of gene expression at post-transcriptional level, yet obtaining quantitative insights into RBP-RNA interactions *in vivo* remains a challenge. Here we developed RBPscan, a method that integrates RNA editing with massively parallel reporter assays (MPRAs) to profile RBP binding *in vivo*. RBPscan fuses the catalytic domain of ADAR to the RBP of interest, using RNA editing of a recorder mRNA as a readout of binding events. We demonstrate its utility in zebrafish embryos, human cells, and yeast, where it quantifies binding strength, resolves dissociation constants, identifies high-specificity motifs for a variety of RBPs, and links binding affinities to their impact on mRNA stability. RBPscan also provides positional information of conserved and novel Pumilio-binding sites in lncRNA *NORAD*. With its simplicity, scalability, and compatibility across systems, RBPscan offers a versatile tool for investigating RBP-RNA interactions and complements established methods for studying post-transcriptional regulatory networks.

**GRAPHICAL ABSTARCT:** 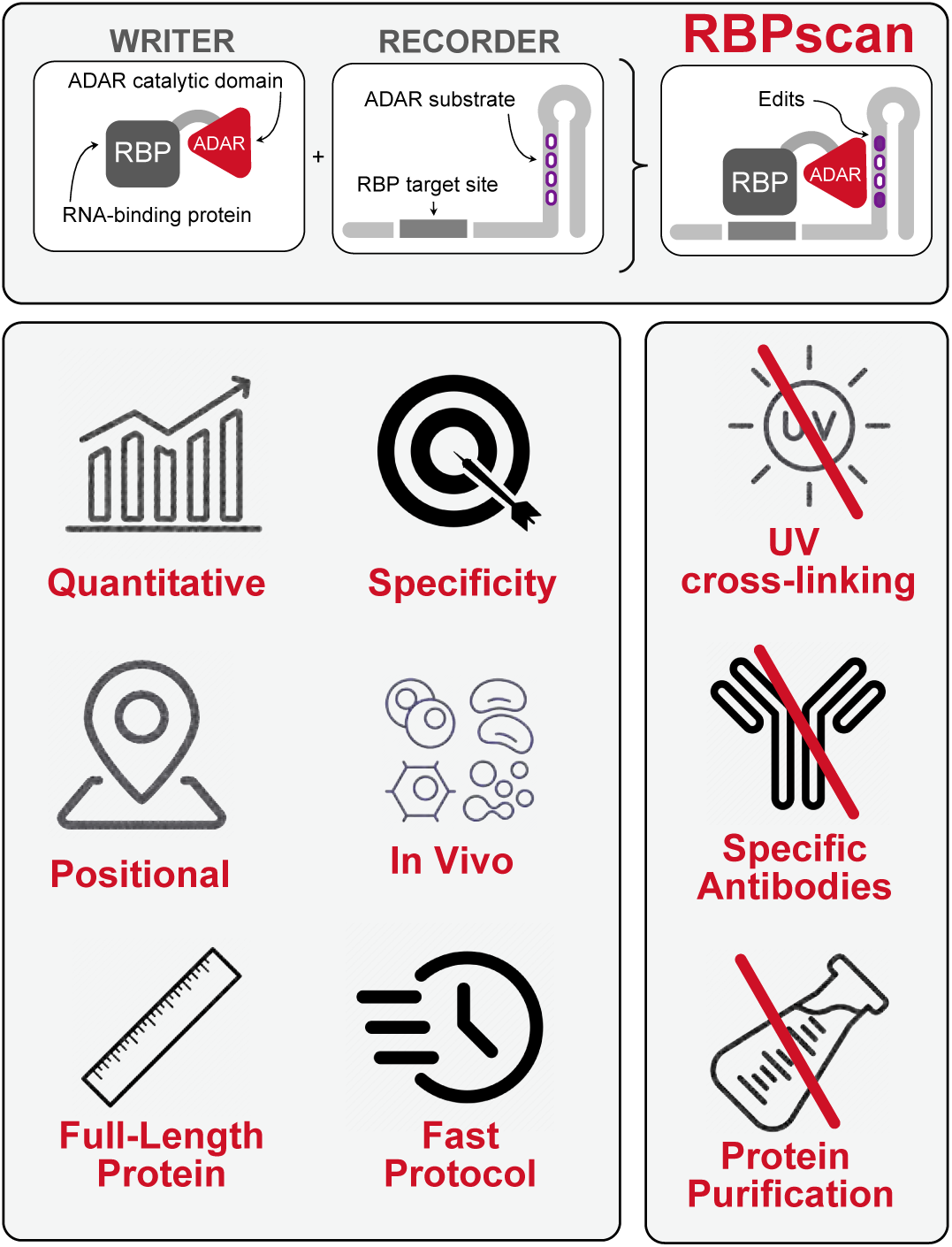

## INTRODUCTION

RNA-binding proteins (RBPs) are essential regulators of post-transcriptional gene expression, orchestrating key steps in an mRNA’s life cycle, from transcription and splicing in the nucleus to translation and decay in the cytoplasm^1^. High-throughput methods such as RNA interactome capture have identified over 1,500 proteins that bind polyadenylated RNA in human cells and other systems, highlighting the vast landscape of RNA–protein interactions^2–5^. RBPs are central to normal development and physiology, and their dysregulation has been implicated in a wide range of diseases, including cancer, neurodegeneration, and developmental disorders^6–8^. Despite their importance, detailed quantitative information about the binding preferences and functional roles of many eukaryotic RBPs remains incomplete.

A variety of methods have been developed to study RBP binding to RNA, each with specific strengths and applications^9^. CLIP-based techniques such as PAR-CLIP, iCLIP, and eCLIP remain invaluable for mapping RBP binding sites *in vivo*, providing positional information about RBP interactions across the transcriptome^10–13^. However, these methods require UV crosslinking, high-quality antibodies, and extensive library preparation, which may limit their scalability^14^. Additionally, CLIP methods often yield qualitative rather than quantitative data, leaving questions about binding dynamics and affinity unanswered.

*In vitro* approaches, including RNA Bind-n-Seq, RNACompete, RNAMaP, SEQRS, and HTR-SELEX, complement CLIP methods by offering quantitative insights into RBP binding preferences^15–19^. These techniques allow for high-resolution characterization of RNA-binding specificities using purified proteins and synthetic RNA libraries. However, they are performed outside the cellular environment, which can affect RBP activity, and typically rely on isolated RNA-binding domains rather than full-length proteins.

More recently, enzyme-based methods such as TRIBE, DART-seq, STAMP, and RNA tagging have expanded the toolkit for studying RBPs^20–23^. These approaches harness RNA-modifying enzymes to mark nucleotides near RBP binding sites *in vivo*. For example, TRIBE uses the catalytic domain of ADAR (adenosine deaminase acting on RNA), while DART-seq and STAMP employ APOBEC, and RNA tagging utilizes Poly(U) polymerase. These tools are relatively straightforward to implement and allow for *in vivo* studies of RBP binding, though the intrinsic sequence preferences of the RNA-modifying enzymes can introduce context-dependent biases that influence interpretation.

RBPscan builds on these advances by combining RNA editing with the scalability of massively parallel reporter assays (MPRAs). In RBPscan, the RBP of interest is fused to the catalytic domain of ADAR, enabling RNA editing as a quantitative readout of RBP–RNA interactions. Synthetic reporter constructs are designed with RNA motifs of interest inserted into the 3’ untranslated region (UTR) alongside a hairpin containing multiple optimal ADAR editing sites. Upon RBP binding, the proximity of ADAR induces targeted adenosine-to-inosine editing, which is detected as A-to-G transitions during sequencing. This straightforward design enables RBPscan to provide quantitative, high-resolution insights into protein–RNA interactions *in vivo* while maintaining simplicity and ease of data analysis. While RBPscan relies on the delivery of exogenous recorders, its broad applicability and adaptability make it a powerful addition to the toolkit for studying RBPs.

## RESULTS

### RBPscan Provides Quantitative, *In Vivo* Insights into Protein–RNA Interactions

To quantitatively determine protein–RNA interactions *in vivo*, independently of UV-crosslinking biases or antibody-based approaches, we developed a dual-reporter system termed **R**NA-**B**inding **P**rotein **S**pecificity and **C**ontextual **A**nalysis via **N**ucleotide editing, or RBPscan. This system comprises a “writer” and a “recorder” (Figure 1A). The writer is a fusion of the RNA-binding protein (RBP) of interest with the catalytic domain of ADAR (ADAR^CD^), while the recorder is a reporter mRNA engineered to include putative binding sites for the RBP along with the Recorder hairpin containing multiple adenosines positioned in perfect conformation for ADAR editing^24,25^ (Figure 1B and S1A). Upon binding of the writer to the recorder mRNA, the proximity of ADAR to its substrates enables deamination of adenosines to inosines, which are read as guanosines during sequencing (Figure S1B). Thus, the number of A-to-G transition mutations serve as a quantitative proxy for protein–RNA interaction strength. To validate RBPscan, we first assessed its specificity using the high-affinity lambdaN22 peptide (λN22) and BoxB interaction pair^26^. One-cell stage zebrafish embryos were injected with various combinations of writer and recorder constructs. Zebrafish embryos offer unique advantages, including the ability to precisely control recorder mRNA amounts via microinjection, thereby circumventing confounding effects from *de novo* transcription or partial delivery systems like lipid-based transfections of cell cultures. Control injections of the BoxB recorder alone or co-injection with either ADAR^CD^ or λN22-ADAR^CD^ in the absence of cognate binding sites (Figure 1C) resulted in minimal background editing (<3%). In contrast, co-injection of λN22-ADAR^CD^ and the BoxB recorder yielded robust editing, with up to 60% of editable sites being modified (Figure 1D). Both Sanger and Illumina sequencing confirmed these findings (Figure 1D and S1C-F). Illumina sequencing revealed a preferential editing hierarchy within the recorder hairpin (Figure S1D). Additionally, we tested that hyperactive ADAR (ADAR^E488Q^)^27^ improved the editing efficiency of the recorder but at the cost of increased editing of the negative controls (Figure S1G). These results demonstrate that endogenous ADAR activity has minimal interference with the recorder system and that functional interactions between the writer and recorder are required for significant editing.

**Figure 1.**
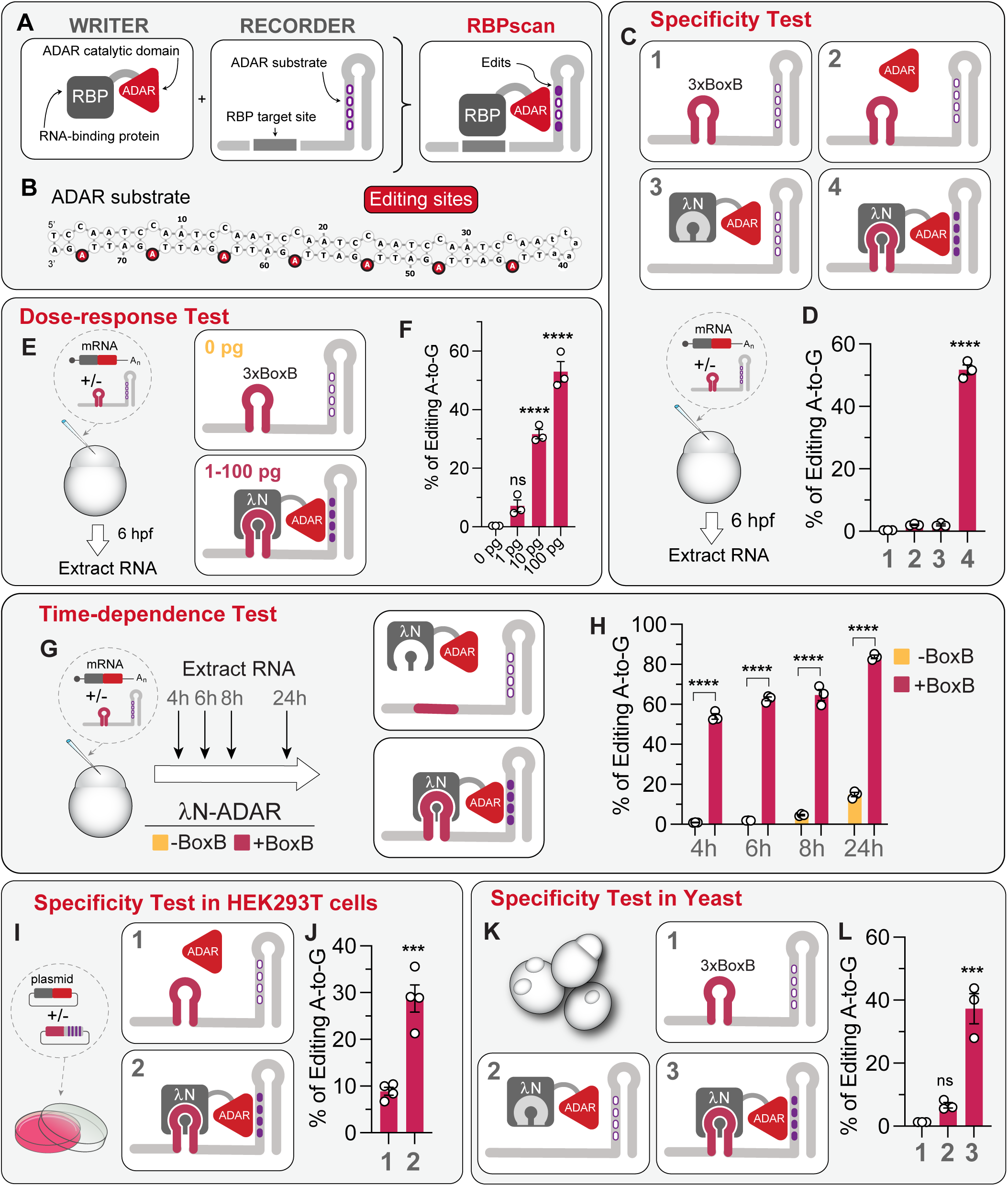
**A)** Cartoon illustrating the components of RBPscan: the “writer” is a RBP of interest fused to the catalytic domain of ADAR. The “recorder” is an mRNA carrying a putative binding site for the RBP and ADAR editing substrate. **B)** Sequence and structure of the ADAR substrate hairpin present in the recorder mRNA. **C)** Cartoon describing the different combinations of writers and recorders to establish robust controls to test the specificity of RBPscan in the experiments conducted in zebrafish embryos upon injection of mRNAs. **D)** Bar plot indicating the % of editing triggered by ADAR^CD^ or λN22-ADAR^CD^ fusion in the corresponding recorders. Wild-type catalytic domain of zebrafish ADAR1 was used. Recorder edits are inferred by Illumina sequencing. Error bars indicate □standard error of the mean of three technical replicates. White circles indicate the mean of individual replicates processed from pools of 30 zebrafish embryos each. *P* value from one-way ANOVA test with Dunnet’s multiple comparison test correction, \*\*\*\**p*<0.0001. **E)** Cartoon describing the experimental design to test the dose-response range of RBPscan in zebrafish embryos. **F)** Bar plot indicating the % of editing triggered by ADAR^CD^ or λN22-ADAR^CD^ fusion in the BoxB recorder. Recorder edits are inferred by Illumina sequencing. Wild-type catalytic domain of zebrafish ADAR1 was used. Values for 0 pg of λN22-ADAR^CD^ (BoxB recorder alone) are reused from Figure 1D for the same condition. Error bars indicate □standard error of the mean of three technical replicates. White circles indicate the mean of individual replicates processed from pools of 30 zebrafish embryos each. *P* value from one-way ANOVA test with Sidak’s multiple comparison test correction, \*\*\*\**p*<0.0001. **G)** Cartoon describing the experimental design to test the time-dependence of RBPscan in zebrafish embryos. **H)** Bar plot indicating the % of editing triggered by λN22-ADAR^CD^ fusion in the empty or BoxB recorder at different time points after embryo fertilization. Wild-type catalytic domain of zebrafish ADAR1 was used. Recorder edits are inferred by Illumina sequencing. Error bars indicate □standard error of the mean of four technical replicates. White circles indicate the mean of individual replicates. *P* value from two-way ANOVA test with Dunnet’s multiple comparison test correction, \*\*\*\**p*<0.0001. **I)** Cartoon illustrating how RBPscan specificity is tested in human HEK293T cells. **J)** Bar plot indicating the % of editing triggered by ADAR^CD^ or λN22-ADAR^CD^ fusion in the BoxB recorder, 24 hours after transfection in human HEK293T cells. The T375G/E488Q mutant of catalytic domain of human ADAR2 was used. Recorder edits are inferred by Sanger. Error bars indicate □standard error of the mean of three technical replicates. White circles indicate the mean of individual replicates. *P* value from an unpaired *t* test. \*\*\**p*=0.0006. Data also shown in Figure S1J. **K)** Cartoon illustrating how RBPscan specificity is tested in human yeast *Saccharomyces cerevisiae*. **L)** Bar plot indicating the % of editing triggered by λN22-ADAR^CD^ fusion in the empty or BoxB recorders, ∼1 day after transformation in yeast. The T375G/E488Q mutant of catalytic domain of human ADAR2 was used. Recorder edits are inferred by Sanger. Error bars indicate □standard error of the mean of four technical replicates. White circles indicate the mean of individual replicates. *P* value from one-way ANOVA test with Dunnet’s multiple comparison test correction, \*\*\**p*<0.0002. Data also shown in Figure S1K.

To evaluate the dynamic range of RBPscan, we conducted dose-response experiments. Zebrafish embryos were injected with a fixed amount of recorder mRNA (100 pg per embryo) and varying doses of writer mRNA (1, 10, or 100 pg per embryo) (Figure 1E). Significant recorder editing was detected at all writer doses, including the lowest tested (1 pg) (Figure 1F and S1H), highlighting the sensitivity of RBPscan over a dynamic range spanning at least two orders of magnitude.

Next, we explored the temporal resolution of RBPscan by analyzing editing at different developmental time points (4, 6, 8, and 24 hours post-fertilization, hpf) (Figure 1G). Significant editing of the BoxB recorder was observed as early as 4 hpf, with editing levels exceeding those of the control by 50-fold. Although background editing increased over time, a significant signal-to-noise ratio (∼5-fold) was maintained at 24 hpf, underscoring the temporal utility of RBPscan (Figure 1H and S1I).

To test versatility of RBPscan, we applied it in human cells and yeast. Plasmids encoding the different ADAR^CD^ variants either with λN22 or without were co-transfected along with BoxB recorder into HEK293T cells (Figure 1I and S1J). Wild-type ADAR^CD^ exhibited higher editing levels but lower signal-to-noise ratios compared to an ADAR^CD^ mutant (T375G/E488Q)^28^ (Figure S1J). Significant signal-to-noise ratios were achieved at 24 hours post-transfection with the ADAR^CD^ T375G/E488Q variant (Figure 1J and S1J). In yeast, we similarly tested the λN22–BoxB pair with both zebrafish and human ADAR^CD^ variants (Figure 1K and S1K). We detected no endogenous background editing, in accordance with absence of ADARs in *S. cerevisiae*. Wild-type zebrafish and human ADAR variants induced strong editing of both the BoxB recorder and the control recorder. Notably, the human ADAR^CD^ T375G/E488Q mutant provided superior signal-to-noise ratios despite lower overall editing efficiency (Figure 1L and S1K).

To enhance the scalability of RBPscan, we developed a fluorescent readout as an alternative to sequencing. In this system, the ADAR editing substrate is encoded within a stop codon (UAG), which, upon editing, is converted to UGG, encoding tryptophan (Figure S2A). The stop codon is strategically positioned between two fluorescent reporter genes, such that editing of the recorder mRNA enables the translation of both reporters, producing a dual-color signal. We validated this approach using recorder mRNAs containing one or two stop codons in zebrafish embryos and HEK293T cells. In the presence of λN22-ADAR^CD^ and the BoxB binding site, we observed a significant increase in green fluorescence (EYFP) in both systems, indicating successful editing and translation of the second reporter (Figure S2B-E). In contrast, very low fluorescence was detected in the absence of the λN22-ADAR^CD^-BoxB interaction. These results establish the validity of a fluorescent RBPscan readout, offering a convenient and multiplex-compatible alternative to sequencing-based approaches.

In summary, RBPscan enables highly specific detection of protein–RNA interactions across a wide dynamic range of expression levels and temporal windows, using a sequencing or fluorescent readouts. Its applicability spans diverse cellular systems, including human cells and yeast, as well as whole organisms such as zebrafish, demonstrating its broad utility for studying protein–RNA interactions *in vivo*.

### RBPscan Resolves Binding Affinities and Quantitative Dynamics of Protein–RNA Interactions

We demonstrated that protein–RNA interactions like the ones described above trigger an editing readout with RBPscan. To evaluate whether the level of editing reflects the strength of protein– RNA interactions, we benchmarked it against Argonaute2 (Ago2), a protein whose binding preferences and interaction strengths are determined by the associated microRNA^29,30^ (Figure 2A). Zebrafish embryos express miR-430 as the dominant microRNA during early development ^31^ (Figure 2B), enabling Ago2-ADAR^CD^ to edit recorders containing the miR-430 8-mer seed matching site (Figure 2C). Inhibition of endogenous miR-430 transcription substantially reduced editing, which was rescued by exogenous pre-miR-430 injection (Figure 2C). To prevent microRNA-mediated degradation of the edited recorder, we incorporated a blocked polyA tail (Figure S3A). To assess the ability of RBPscan to distinguish between binding affinities of different miR-430 seed variants (8-mer, 7-mers, and 6-mers) (Figure 2B), we introduced a pool of recorders encoding these sites. RBPscan data accurately ranked seed variants according to their binding strength (Figure 2D and Tables S2), in correlation with their capacity to induce mRNA decay as determined by a massively parallel reporter assay^32^ (Figure 2E).

**Figure 2.**
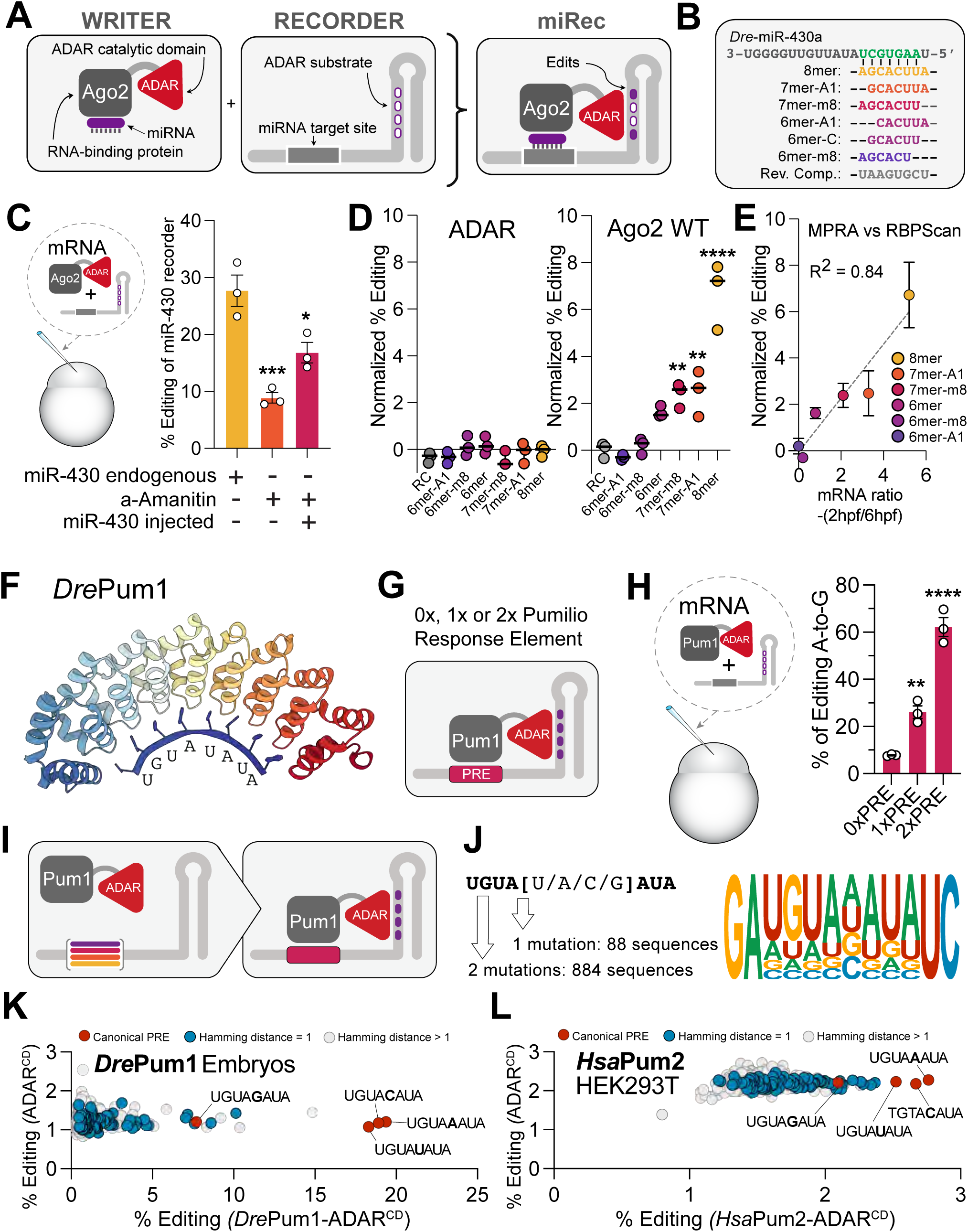
**A)** Cartoon illustrating the writer and recorder design when Ago2 is used as the writer. In this case, RBPscan is testing a ternary interaction between Ago2, the associated miRNA, and the target of the corresponding miRNA. **B)** Seed matching series of miR-430, the most abundant miRNA in early zebrafish embryogenesis. Seeds matches are shown 5’-to-3’, as they would be found in the 3’UTR of a target mRNA. **C)** RBPscan experiment showing how the editing triggered by Ago2-ADAR^CD^ in presence of endogenous miR-430 is abrogated by the inhibition of miR-430 expression (α-amanitin) and rescued by pre-miR-430 injection. Recorder edits are inferred by Sanger sequencing. Catalytic domain of wild-type zebrafish ADAR1 was used. Error bars indicate □standard error of the mean of three technical replicates. White circles indicate the mean of individual replicates processed from 30 zebrafish embryos each. *P* values from one-way ANOVA test, \**p*=0.0139, \*\*\**p*=0.0009. **D)** RBPscan experiment showing the edits triggered by zebrafish ADAR^CD^ or Ago2-ADAR^CD^ on the pooled library containing the miR-430 seed series (Figure 2B). Catalytic domain of wild-type zebrafish ADAR1 was used. Individual values for each technical replicate are shown as colored circles. Horizontal black line indicates the median. *P* values from one-way ANOVA test with Dunnet’s multiple comparison test correction, \*\**p*<0.01, \*\*\*\**p*<0.0001. **E)** Benchmarking of the RBPscan results from Figure D for the miR-430 seed series against actual mRNA decay data triggered by miR-430 from a massively parallel reporter assay^32^. **F)** AlphaFold3 structure prediction of the Pum-HD domain array of zebrafish Pumilio 1 (Pum1), here shown interacting with an 8-nucleotide RNA molecule containing the motif UGUAUAUA. **G)** Cartoon illustrating the strategy to test multiple PRE with zebrafish Pum1-ADAR^CD^ in a RBPscan assay. **H)** Recording of the interaction of Pum1-ADAR^CD^ with a variable number of PREs. Recorder edits are inferred by Sanger sequencing. Error bars indicate □standard error of the mean of three technical replicates. White circles indicate the mean of individual replicates processed from 30 zebrafish embryos each. *P* values from one-way ANOVA test with Dunnet’s multiple comparison test correction, \*\**p*=0.0066, \*\*\*\**p*<0.0001. **I)** A cartoon illustrating the strategy for using the RBPscan assay to screen the binding of zebrafish Pum1 to 1,000 PRE variants. Only the motifs recognized by Pum1 will induce editing of the recorder **J)** Sequence logo of the designed PRE recorder library, which includes all motifs differing from the canonical PREs by 1- and 2-nucleotide permutations, along with non-PRE sequences. **K)** Scatter plot showing the % of editing of individual motifs after co-injection into zebrafish embryos together with 10 pg of mRNA encoding ADAR^CD^ or zebrafish Pum1-ADAR^CD^. Catalytic domain of wild-type zebrafish ADAR1 was used. Each circle represents the mean editing of a given motif in three replicates. Circles are colored according to their Hamming distance to a canonical PRE. **L)** Scatter plot showing the % of editing of individual motifs after transfection of HEK293T cells together with ADAR^CD^ or human Pum2-ADAR^CD^. Catalytic domain of wild-type human ADAR2 was used. Each circle represents the mean editing of a given motif in three replicates. Circles are colored according to their Hamming distance to a canonical PRE.

To further evaluate the quantitative capabilities of RBPscan, we employed the full length RNA-binding protein Pumilio (Pum1) (Figure 2F), which binds a well-characterized consensus motif UGUAHAUA, the Pumilio Response Element (PRE)^33–35^. Recorders containing 0, 1, or 2 PRE sites were tested with Pum1-ADAR^CD^ (Figure 2G). Editing levels were nearly doubled in recorders with 2 PREs compared to those with a single site, indicating that the RBPscan editing readout scales with RNA-protein interaction strength (Figure 2H). Adding two binding sites inevitably places one site farther from the recorder hairpin. However, when we tested a series of recorders where the PRE was separated by a spacer sequence of increasing length and depleted of PREs (Figure S2B), we found that a PRE located up to 174 nucleotides away from the recorder hairpin remained equally functional (Figure S2C).

To further assess the resolution of RBPscan, we challenged Pum1-ADAR^CD^ with a library of recorders containing various PRE permutations. (Figure 2I). The PRE library consisted of 1,000 sequences (Table S3), which were designed by systematically introducing all possible single- and double-nucleotide mutations of the consensus PRE motif, along with non-PRE controls (Figure 2J). Following co-injection into zebrafish embryos, the editing landscape was assessed via high-throughput Illumina sequencing, facilitated by partial adaptors flanking the reporter region. While overall editing levels across the library were low (<4%) (Figure S2D), Pum1-ADAR^CD^ robustly edited three specific recorders to levels as high as 18% (Figure 2K). Motif analysis of the highly edited recorders revealed precise matches to the UGUAHAUA consensus. By contrast, motifs with deviations, such as UGUAGAUA (carrying a G at position 5), scored significantly lower (7.5% editing) (Figure 2K).

Importantly, the specificity towards motifs was independent of Pum1-ADAR^CD^ dosage, as similar results were observed with injections of both 10 pg and 100 pg of writer mRNA (Figure S2E and F). Comparable motif segregation was also observed when human Pum2 (Pum2-ADAR^CD^) was transfected in HEK293T cells, as the UGUAHAUA motifs where consistently the top-edited motifs (Figure 2L and S2G). In HEK293T cells, wild-type ADAR outperformed the mutant ADAR (ADAR^CD^ ^T375G/E488Q^) in resolving the PRE recorder library (Figure S2H). Notably, the UGUAHAUA motifs are not merely enriched in the group of motifs that trigger editing, but actually each rank #1, #2, and #3 of the most edited motifs (Table S4), regardless of the experimental system. These results underscore the ability of RBPscan to discover biologically relevant binding sites in different experimental systems and its potential for identifying the binding preferences of RBPs with unknown specificities.

### RBPscan Quantifies Relative Binding Affinities and Links RBP–RNA Interactions to Functional Outcomes

Given that RBPscan can rank motifs based on the affinity for the RBP of interest and detect these interactions across at least two orders of magnitude of RBP expression levels, we decided to leverage these features to perform dose-response experiments for RBPs *in vivo*.

We injected a fixed amount of the PRE mRNA library (100 pg/embryo) described above along with increasing doses (0.9 to 120 pg/embryo) of Pum1-ADAR^CD^ mRNA into one-cell-stage zebrafish embryos. After collecting and analyzing each condition in triplicates, we generated dose-response curves for Pum1 binding to 1000 motifs present in the library (Figure 3A,B,C,D and Table S5). From these sigmoidal curves, we calculated apparent relative dissociation constants (*K_D_*) based on the inflection points, allowing us to rank the motifs by affinity. The results aligned closely with the known binding preferences of Pum1. The canonical UGUAHAUA motifs ranked as the top three with similar apparent *K_D_* values (15.38□2.20, 15.39□2.10, and 15.66□2.45 with standard error), while the UGUAGAUA motif displayed a significantly weaker affinity (18.34□1.60). The units of these *K_D_*, pg of Pum1-ADAR^CD^ mRNA per embryo should be equivalent to Pum1 protein concentration if we assume constant translation and editing rates in the confined volume of the embryo.

**Figure 3.**
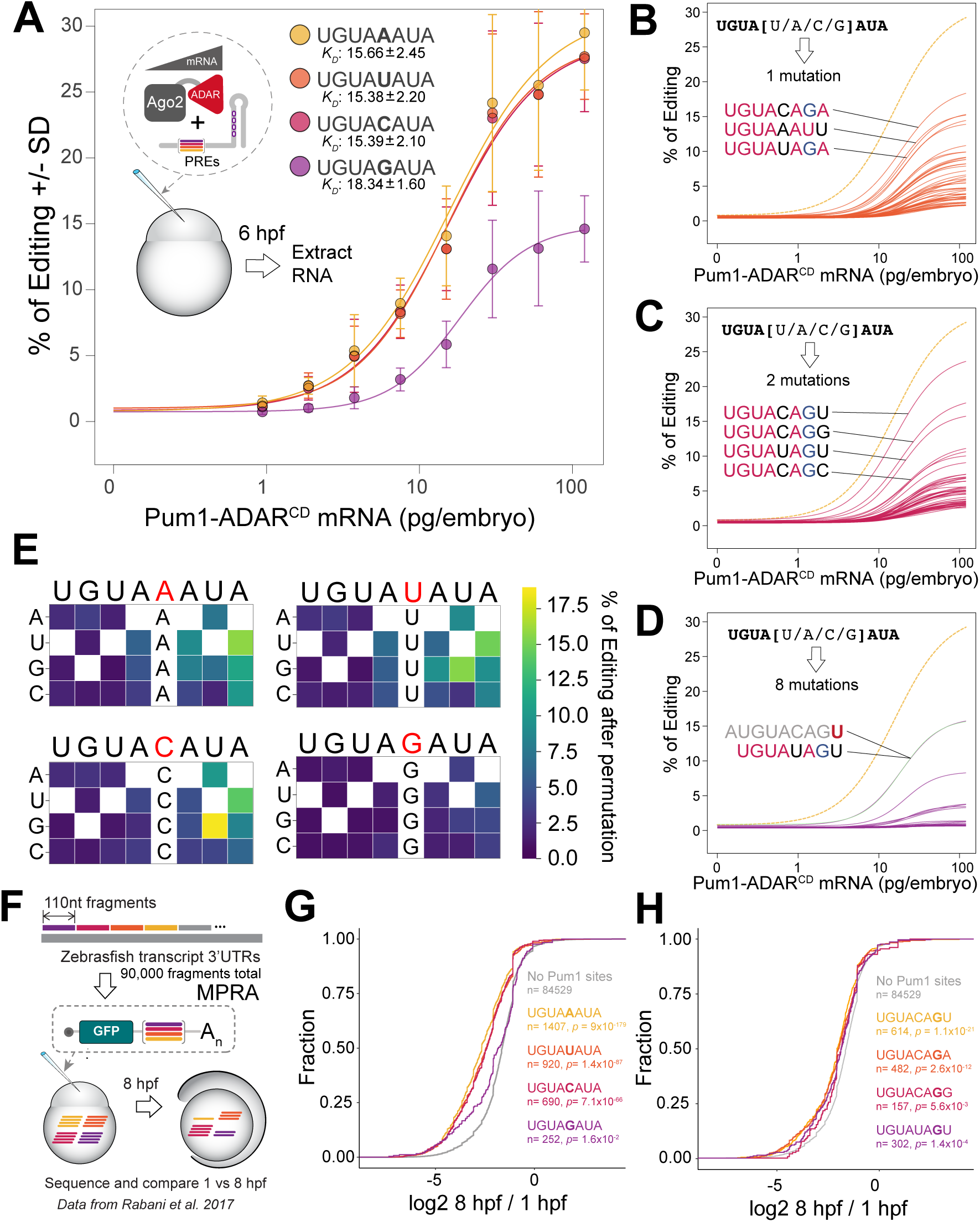
**A)** Dose-response curves of Pum1-ADAR^CD^ mRNA injected at different doses (0.9-120 pg per embryo) against a fixed amount of the PRE variant recorder library. Catalytic domain of wild-type zebrafish ADAR1 was used. The curves represented here are for the canonical Pum1 binding motifs. Error bars indicate standard error of the mean. Colored circles indicate the average value of three triplicate samples. **B)** Similar to Figure 3A, but showing dose-response curves for Pum1 against binding sites that differ from the canonical PREs by a single mutation (Hamming distance = 1). **C)** Same as Figure 3A, but the dose-response curves represented belong to Pum1 against binding sites that differ by 2 mutations from the canonical PRE. **D)** Same as Figure 3A, but the dose-response curves represented belong to Pum1 against binding sites that differ by 8 mutations from the canonical PRE. **E)** Heat map showing Pum1 affinity for motifs that differ by 1 nucleotide from the four canonical PRE. **F)** Cartoon describing the massively parallel reporter assay done in zebrafish embryos by Rabani et al.^32^ to identify regulatory motifs. **G)** Cumulative plot showing the extend of mRNA decay elicited by each of the indicated PREs. **H)** Same as Figure 3G, but showcasing the decay induced by the novel Pumilio binding motifs.

When analyzing the full spectrum of motifs in the library, we observed that variants differing from the consensus by a single mutation (Hamming distance = 1) provided new insights into Pum1 binding mechanisms. Mutations in the first four nucleotides of the motif almost completely abolished Pum1 binding, suggesting these nucleotides serve as a “seed” for recognition (Figure 3B and 3E). In contrast, mutations in the second half of the motif were less disruptive, where a G at position 7 was the most tolerated. Notably, the previously unappreciated motif UGUACAGU, containing a G at position 7 and a U mutation at position 8 (Hamming distance = 2), displayed higher activity than any single-mutation variant, indicating a compounding effect between these positions (Figure 3C).

Further analysis revealed an unexpected high-affinity interaction among one of the control motifs designed to lack functional binding sites (Hamming distance = 8). Inspection of these sequences uncovered a regenerated PRE, shifted by one nucleotide and using an uracil present in 3’ flanking sequence of the cloning site (Figure 3D). The dose-response curves for the shifted motif and the consensus motif overlapped, confirming that editing is exclusively driven by sequences within the recorder construct. Together, these experiments demonstrate that RBPscan can quantify the relative affinities of protein–RNA interactions *in vivo* and establish a robust framework for detailed motif analysis of RBPs.

To benchmark how Pum1 binding affinity influences its functional role in mRNA repression, we re-analyzed data from a massively parallel reporter assay performed in early zebrafish embryos ^32^ (Figure 3F). During early embryogenesis, Pum1 mediates the destabilization and decay of numerous transcripts. We assessed the stability of fragments containing each PRE variant relative to controls lacking PRE sites. The canonical UGUAHAUA motifs induced reporter mRNA decay with similar magnitudes as predicted by their relative *K_D_* values (Figure 3G). The weaker UGUAGAUA motif induced significant but less pronounced decay. Strikingly, the newly discovered UGUACAGU motif and its derivatives also drove robust mRNA decay responses, despite not being previously associated with Pum1 function (Figure 3H). These findings establish a critical link between binding affinity as determined by RBPscan and the functional consequences of RBP–RNA interactions.

These results collectively demonstrate that the output of RBPscan is proportional to the strength of protein–RNA interactions and quantitatively reflects the magnitude of the downstream effects of these interactions. While RBPscan does not provide absolute dissociation constants, its ability to quantify relative apparent *K_D_* values *in vivo* offers powerful advantages. These include: (1) precise analysis of RBP target specificity, (2) predictive modeling of functional outcomes from binding events, and (3) identification of secondary binding motifs with functional relevance.

### RBPscan Identifies Binding Motifs for Diverse RBPs with High Specificity and Versatility

For RBPscan to become a widely applicable tool for studying protein–RNA interactions, it must be adaptable to a broad repertoire of RBPs. To test its versatility, we sought to identify the binding motifs of diverse RBPs using a library of recorders containing randomized 7-mer binding sites, encompassing all 16,384 possible motifs. To accommodate RBPs with 8-mer motifs, such as Pumilio and Argonaute, we fixed the nucleotide immediately following the 7N sequence as an A, an integral part of their canonical motifs.

We first validated this expanded recorder library with Ago2-ADAR^CD^. Triplicate library screenings revealed that the most enriched motifs corresponded to the miR-430 seed series—the dominant miRNA expressed in early zebrafish embryos—with the 8-mer motif exhibiting the highest editing levels (Figure 4) and triggering the largest decay of miR-430 targets (Figure S4A). As a control, library screening in embryos with inhibited miR-430 expression by α-amanitin yielded significantly fewer edited motifs, edited at lower levels, none of which matched any miR-430 seed, confirming specificity.

**Figure 4.**
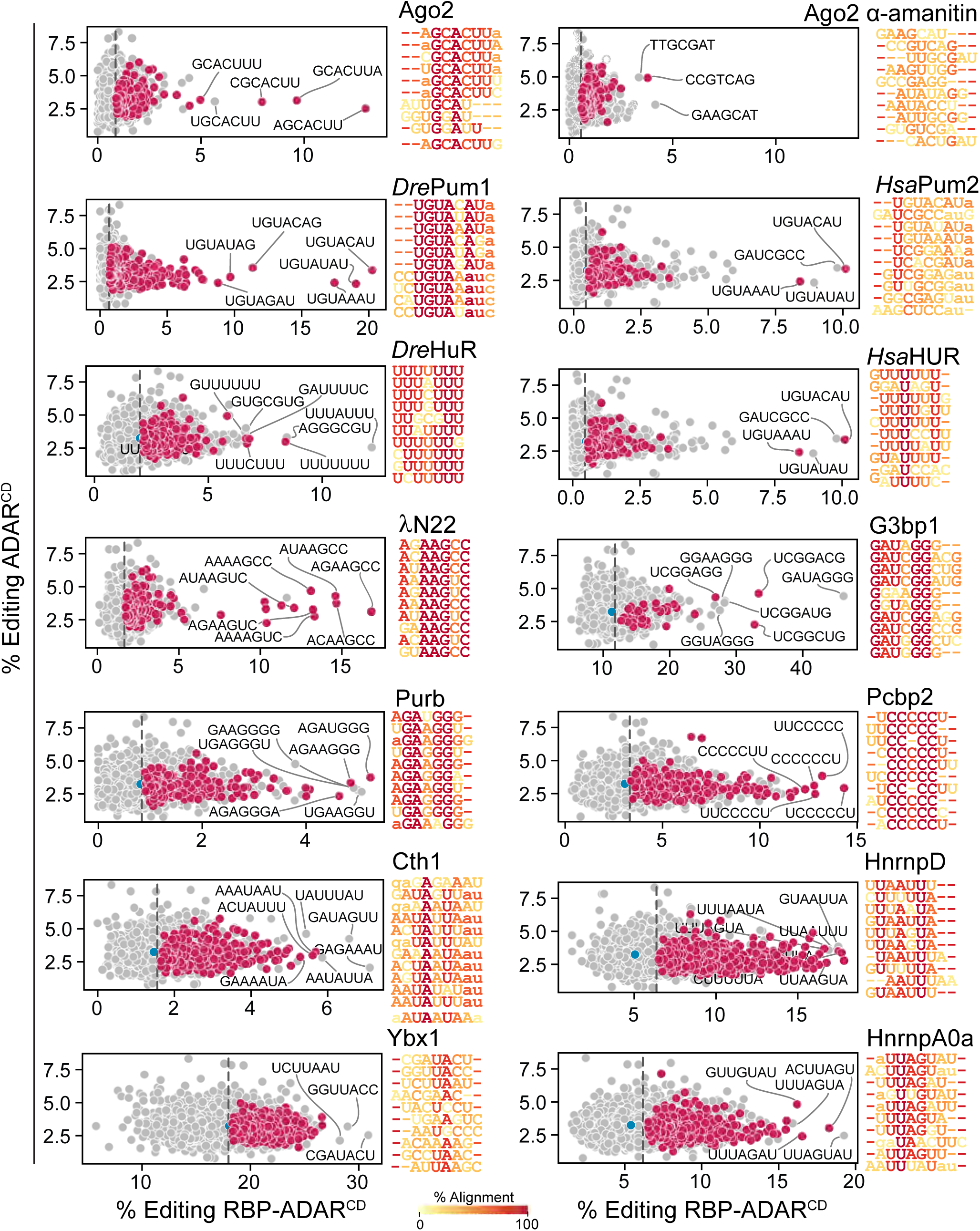
**A)** Scatter plots showing the editing levels of a library of recorders containing all possible 7-mer motifs (16,384) when co-injected into zebrafish embryos with ADAR^CD^ or the ADAR^CD^ fusion with the corresponding RBP. Catalytic domain of wild-type zebrafish ADAR1 was used. Motifs colored in red display editing levels 2 standard deviation above the average editing across all motifs. Blue dot corresponds to the empty recorder without any 7-mer motif. Next to the scatter plot we show the alignment of the topmost edited motifs (top to bottom) for each RBP. Nucleotides are colored according to their prevalence in the alignment position. Lower cases indicate the nucleotides present in the fixed positions adjacent to the cloned motif.

Next, we tested zebrafish Pum1 and human Pum2. For Pum1, the top 10 edited motifs all included the UGUA seed, consistent with our results from the smaller library. Human Pum2-ADAR^CD^ similarly identified motifs matching its consensus sequence, although the top 10 motifs included additional mixed sequences. We extended our analysis to HuR, testing both zebrafish and human orthologs. For both, U-rich motifs dominated the top-edited sequences (Figure 4), in agreement with known consensus sequences for HuR^36^.

To explore whether RBPscan can study structured RNA-binding sites in addition to linear motifs, we screened for motifs interacting with the λN22 peptide, which recognizes the structured BoxB hairpin. Despite using a randomized 7-mer library incapable of forming hairpins on its own, we identified high-affinity binding motifs. Further investigation revealed that these motifs, in conjunction with surrounding flanking sequences, reconstituted a hairpin structure nearly identical to BoxB (Figure S4B). Notably, position 2 of the 7-mer motif tolerated any nucleotide, consistent with structural data on (λN22–BoxB interactions that demonstrated that the nucleotide in this position does not participate in the stacking interaction^37^. In addition, we observed a transition from a G:C pairing to G:U wobble in the stem, which are tolerated as either type of pairing will retain the hairpin structure, These results demonstrate RBPscan’s capability to study both linear and structured RNA-binding motifs.

A particularly notable case was Ybx1, which is not thought to bind RNA in a sequence-specific manner. Instead, RBPscan revealed uniformly elevated editing across all motifs (Figure 4), suggesting that Ybx1 interacts with the recorder mRNAs regardless of motif sequence^38^.

We continued identifying top-edited motifs for additional RBP–ADAR^CD^ fusions, including G3bp1, Purb, Pcbp2, Cth1, HnrnpD, and HnrnpA0 (Figure 4). For all proteins, the most edited motifs matched or closely resembled their consensus motifs determined by established techniques such as SELEX, CLIP, or Bind-n-Seq. Indeed, we benchmarked our motif discovery for Pum1, HnrnpA0a, HnrnpD, and Pcbp2 against motifs enriched *in vitro* for their human orthologs using RNA Bind-n-Seq^39^, a technique with two key differences from RBPscan: (1) it is performed *in vitro*, and (2) it uses truncated proteins to facilitate purification. Despite these differences, we observed that the motifs identified by both methods showed stronger correlation at higher doses of RBP in the Bind-n-Seq assay (Figure S5 and Table S7). These findings demonstrate that RBPscan reliably identifies *bona fide* binding motifs of RBPs *in vivo*, providing complementary insights to *in vitro* approaches.

For each tested RBP, we provide the sequences of the ten most edited motifs along with their associated activity scores (see Table S6 for complete list of motifs). Importantly, we deliberately avoided collapsing these motifs into a single consensus sequence. While traditional sequence logos highlight conserved positions, they often fail to capture the contextual interdependence between nucleotides in adjacent positions. To demonstrate that such context matters and outperforms random connections in a consensus sequence, we applied machine learning to identify motifs with the greatest impact on mRNA decay. We analyzed four RBPs with established roles in destabilizing mRNAs: Pum1, Ago2, Pcbp2, and Cth1. For Pum1, Ago2, and Pcbp2, motif rankings based on editing induction correlated strongly with inferred decay capacity (Pearson Correlation Coefficent R = 0.976, 0.622, and 0.739, respectively) (Figure S4C). Motifs ranked at the bottom of the list showed minimal predicted impact, underscoring the importance of curated motif lists with associated activity scores over generalized consensus sequences for accurately predicting post-transcriptional regulatory potential. In contrast, Cth1 motifs showed no correlation with the decay predictions. While the identified motifs were AU-rich, consistent with Cth1 binding preferences, we hypothesize that Cth1 may be inactive in the context of the MPRA experiment or that our library did not sufficiently enrich for functional sequences in the top ten motifs.

Altogether, these results showcase the potential of RBPscan to systematically uncover linear and structural binding motifs for a wide range of RBPs. The assay’s high specificity eliminates the need for prior knowledge, as motifs with the highest affinity—as confirmed by orthogonal techniques—consistently emerge as the most edited sequences.

### RBPscan Accurately Identifies the Location of RBP Target Sites in Transcripts

A critical step in elucidating post-transcriptional regulatory networks is identifying the mRNA targets of a given RBP. Techniques such as CLIP, TRIBE, and the more recent STAMP, infer RBP binding sites by performing peak analyses over clustered reads. To evaluate RBPscan’s capability to accurately delineate RBP target sites in transcripts, we analyzed Pumilio binding sites in *NORAD*, a long non-coding RNA (∼5 kb) containing multiple Pumilio binding sites critical for regulating genome stability in mammals^40,41^.

To uncover Pumilio binding sites in *NORAD* using RBPscan, we designed a tiling library of recorders, each encoding 30-nucleotide fragments of human *NORAD* with a 5-nucleotide displacement between overlapping fragments (Figure 5A). This comprehensive library comprised more than 1,070 unique recorders (Table S8), ensuring high-resolution coverage. We injected this library into zebrafish embryos, alone or co-injected with ADAR^CD^ or Pum-ADAR^CD^. In controls with the library alone or with ADAR^CD^, we observed uniform editing across all nucleotides in *NORAD* (Figure 5B), independent of sequence or position, with slightly elevated editing in the presence of ADAR^CD^ (Figure 5C). However, co-injection of *Dre*Pum1-ADAR^CD^ (Figure 5D) or *Hsa*Pum2-ADAR^CD^ (Figure 5E) resulted in highly localized editing peaks, up to ∼100-fold higher than background levels. These peaks precisely aligned with previously annotated canonical Pum1 binding sites. Further analysis revealed additional functional Pum1 binding sites under smaller peaks detected by RBPscan (Figure S5A-B), increasing the total number of Pumilio binding sites in *NORAD* from 15 to 35.

**Figure 5.**
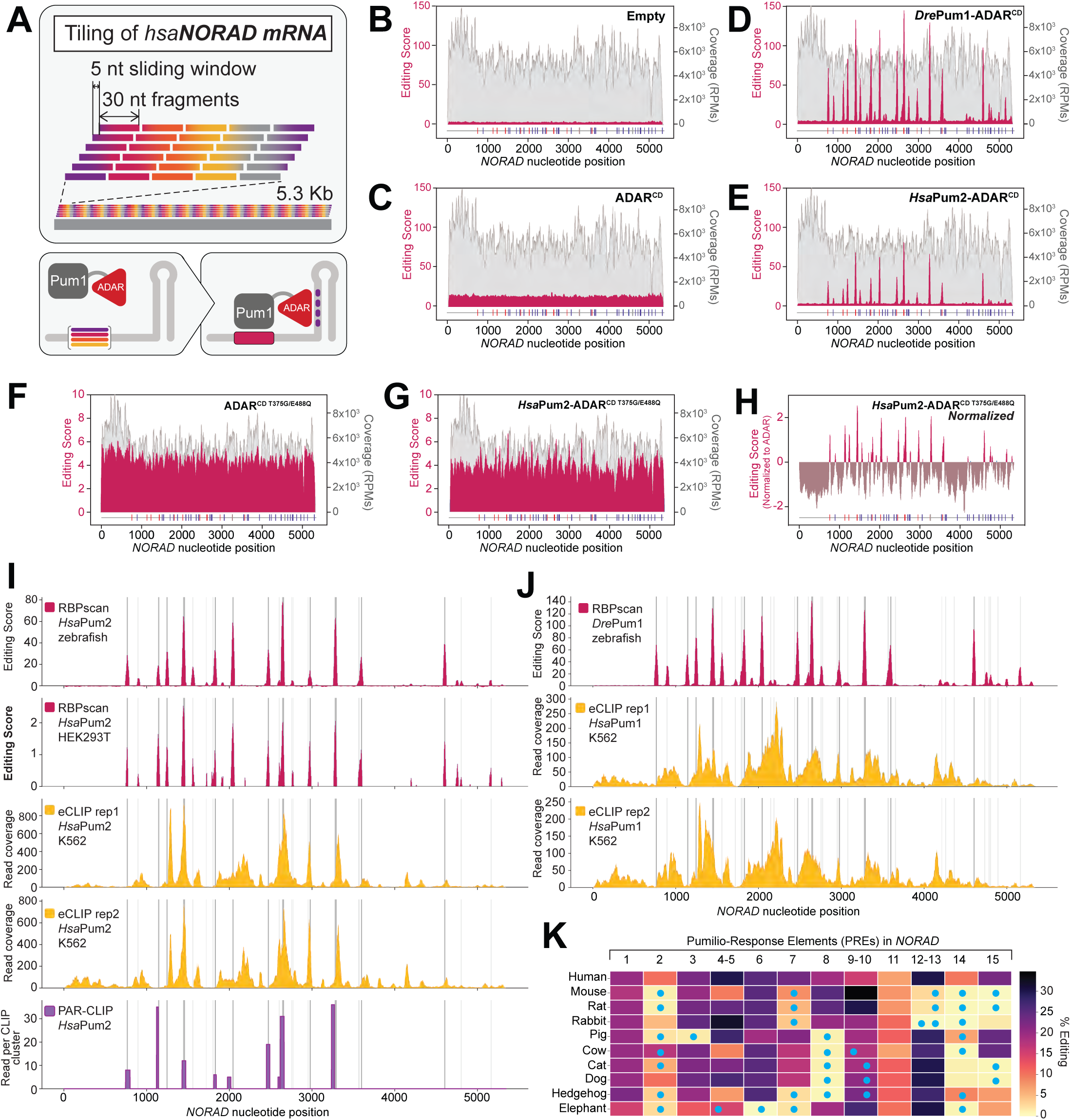
**A)** Cartoon describing the tilling strategy to analyze the binding of Pumilio proteins to human *NORAD RNA* with RBPscan. **B)** Recorder read coverage (grey) and editing score (red) after injection of the *NORAD* tilling recorder library in zebrafish embryos without any writer. **C)** Recorder read coverage (grey) and editing score (red) after injection of the *NORAD* tilling recorder library in zebrafish embryos with ADAR^CD^. **D)** Recorder read coverage (grey) and editing score (red) after injection of the *NORAD* tilling recorder library in zebrafish embryos with *Dre*Pum1-ADAR^CD^. **E)** Recorder read coverage (grey) and editing score (red) after injection of the *NORAD* tilling recorder library in zebrafish embryos with *Hsa*Pum2-ADAR^CD^. **F)** Recorder read coverage (grey) and editing score (red) after transfection of the *NORAD* tilling recorder library in HEK293T cells with ADAR^CD^ ^T375G/E488Q^. **G)** Recorder read coverage (grey) and editing score (red) after transfection of the *NORAD* tilling recorder library in HEK293T cells with *Hsa*Pum2ADAR^CD^ ^T375G/E488Q^. **H)** Editing score (red) triggered by *Hsa*Pum2ADAR^CD^ ^T375G/E488Q^ after normalization by ADAR^CD T375G/E488Q^ editing. **I)** Gene browser tracks showing the estimated location of Pum2 binding sites inferred by RBPscan and normalized by background editing, eCLIP, or PAR-CLIP. Thick vertical grey lines indicate the location of canonical Pumilio response elements, thin grey lines indicate the position of other sites that at least contain the UGUA seed of the Pumilio binding site and RBPscan score above 4 (*dre*Pum1) or 3 (*hsa*Pum2). **J)** Same as Figure 5I, but for *Dre*Pum1 normalized by background editing. **K)** Heat map showing the editing triggered by Pumilio response elements of different species at the 15 main positions described initially in Lee at al^40^. Blue dots indicate sequences that after alignment to human *NORAD* do not show the core UGUA in the same position.

Given the accessibility of cell culture systems to most RNA biology laboratories, we tested RBPscan’s performance in identifying human Pum2 binding sites in *NORAD* using HEK293T cells. Transfection of both Pum2-ADAR^CD^ and the *NORAD mRNA library* produced editing profiles similar to those observed from ADAR^CD^ alone (Figure 5F-G). However, after normalizing Pum2-ADAR^CD^ edits against ADAR^CD^ control levels, a clear pattern of peaks emerged (Figure 5H), precisely overlapping with annotated sites and those uncovered in zebrafish, underscoring the reproducibility and platform-independence of RBPscan.

We next benchmarked RBPscan against established tools such as PAR-CLIP and eCLIP. PAR-CLIP identified 9 of the 15 canonical PREs annotated in *NORAD* for *Hsa*Pum2 (Figure 5I), while RBPscan not only detected all 15 but also uncovered an additional 7 for *Hsa*Pum2 and 20 functional sites for *Dre*Pum1 (Figure 5J and S5B). Comparison with eCLIP revealed narrower RBPscan peaks, with an average width of 15 nucleotides compared to the broader, near-continuous read clusters typical of eCLIP. Furthermore, eCLIP peaks are inherently biased to lie downstream of RBP binding sites due to protocol constrains^11^, whereas RBPscan peaks precisely overlay the binding motif itself, simplifying motif identification and analysis.

Notably, RBPscan results were consistent across experimental platforms, whether in zebrafish embryos or HEK293T cells, indicating that Pumilio binding affinity is similar between othologues^42,43^ and independent of cellular contexts. The binding affinities of Pumilio sites in *NORAD* also showed remarkable conservation across species. When we examined the conservation of the sequence and binding strength of Pumilio sites in *NORAD* across mammalian species, we found that if the site was present, binding activity remained within range of the strength of the top motif (Figure 5K). These findings suggest that evolutionary pressure conserves the functional strength of Pumilio interactions at precise locations rather than the sequence of their target sites.

In summary, these results demonstrate RBPscan’s ability to pinpoint the position of RBP binding sites in transcripts with high accuracy and specificity. Future applications of RBPscan using complex libraries derived from random transcriptome fragments hold the potential to enable transcriptome-wide discovery of RBP targets.

### Limitations of the Technique

A key limitation of RBPscan is that, while it measures RBP–RNA interactions in their native cellular context, both the RBP and the target RNA must be transduced and overexpressed in the experimental system as a protein fusion with ADAR^CD^. As a result, the use of RBPscan is restricted to systems amenable to genetic manipulation, whether transient (e.g., transfection or microinjection) or stable (e.g., lentiviral transduction or transgenesis). Additionally, unlike *in vitro* techniques, RBPscan is subject to potential confounding effects from RBP complexes. If the RBP of interest forms a complex with other RBPs, the extracted motifs may represent a composite of the binding preferences of the complex rather than the isolated RBP. This interplay underscores the importance of carefully interpreting results in the context of the broader cellular environment.

## DISCUSSION

Accurate identification and quantification of protein–RNA interactions are critical for decoding post-transcriptional regulatory networks. In this study, we developed RBPscan, a novel tool that integrates RNA editing with massively parallel reporter assays (MPRAs). RBPscan provides quantitative measurements of protein–RNA interaction strength, identification of binding motifs, affinity resolution through dose-response analyses, and positional information of interactions. Importantly, RBPscan is versatile and can be applied across diverse platforms, including human cell lines, yeast, and zebrafish.

RBPscan complements the current toolkit for studying RNA–protein interactions. Unlike existing methods, RBPscan provides quantitative data *in vivo* using full-length proteins, without relying on protein purification, antibody availability, or UV crosslinking. The use of standardized reporter constructs ensures reliable editing that is independent of the distribution of endogenous editing substrates in target RNAs. RBPscan builds directly on approaches such as TRIBE^20^ and STAMP^22^ yet addresses a key limitation of these methods: the reliance on non-uniform endogenous substrates for the corresponding RNA editing enzymes, which reduces binding site resolution. By introducing a separate reporter mRNA encoding multiple perfect ADAR substrates, RBPscan uncouples the editing process from endogenous RNA contexts, enabling precise quantification of binding events.

Although we benchmarked RBPscan with proteins of known interactions, it uncovered novel high-affinity binding sites for Pumilio. These sites deviate significantly from the canonical consensus motif but still induce Pumilio-mediated RNA decay *in vivo*. Furthermore, RBPscan resolved the positions of previously annotated Pumilio binding sites in *NORAD* and identified 20 additional high-affinity sites that were undetectable using standard techniques due to resolution constraints.

The versatility of RBPscan makes it particularly suitable for specific applications. These include profiling binding motifs of novel RNA-binding proteins with unknown specificities, investigating the contribution of RNA structure to RBP binding affinity, and performing transcriptome-wide profiling of RBP binding sites using random RNA fragment libraries. By simultaneously resolving the strength and position of RBP interactions within transcripts, RBPscan enables deeper insights into protein–RNA interaction dynamics. For example, our analysis of Pumilio interactions with *NORAD* revealed that the conservation of binding sites across species correlates with motif strength and position, rather than the precise sequence of the motif. Extending this approach to other RBPs could significantly advance our understanding of post-transcriptional regulatory networks.

The current application of RBPscan is limited to systems amenable to transfection, injection, transgenesis, or viral transduction, as both writer and recorder constructs must be introduced into cells. Future developments include lentiviral delivery systems to enable broader applicability across cell types and tissues, as well as increasing resolution to analyze RBPscan at single-cell levels.

In summary, RBPscan combines simplicity, versatility, and quantitative power, making it a transformative tool for studying RNA–protein interactions *in vivo*. By offering direct *in vivo* quantification and positional information for full-length proteins, RBPscan complements existing approaches while addressing longstanding challenges. The insights provided by RBPscan have the potential to fundamentally advance our understanding of post-transcriptional gene regulation and to drive new discoveries in RNA biology.

## Supporting information

Table_S1_oligonucleotides

Table_S2_miR-430_pool

Table_S3_PRE_oligos

Table_S4_realated to figure 2

Table_S5_Related_to_Figure3

Table_S6_RBPscan_All_RBPs_Related_to_Figure4

Table_S7_Bind-n-Seq

Table_S8_NORAD_oligos

Table_S9_NORAD_RBPscan

## Resource availability

Plasmids will be deposited in the plasmid repository Addgene. Illumina sequencing data will be deposited at the Sequence Read Archive (SRA). Code used for the analysis of the data will be available in GitHub.

## Acknowledgments

We thank M. Garcia-Marcos, and A. Grishok for fruitful discussions of the project and access to instruments and microscopes. This project was funded with an Ignition Award grant from Boston University to DC and an Early Career Development award to DK from the Department of Biochemistry and Cell Biology. DK is grateful to the organizers of CSHL “Programming for Biology” course.

## Author contributions

D.A.K. designed, performed, and analyzed all the experiments and helped with the writing of the manuscript. O.S. helped with the Ago2 experiments and Bind-n-Seq comparisons, T.M. helped with cell culture experiments, E.P. helped with plasmid cloning. I. conducted the RBPscan experiments in yeast. S.W. Conducted the machine learning experiments, D.C. conceived and supervised the project, performed experiments, analyzed the data, acquired funding, assembled the figures, and wrote the manuscript.

## Supplemental information

Supplementary Tables ST1: List of oligonucleotides.

Supplementary Tables ST2: Data relevant to Figure 2.

Supplementary Tables ST3: List of oligonucleotides used to clone the PRE library.

Supplementary Tables ST4: Data relevant to Figure 2.

Supplementary Tables ST5: Data relevant to Figure 3.

Supplementary Tables ST6: Data relevant to Figure 4.

Supplementary Tables ST7: Data relevant to Figure S5.

Supplementary Tables ST8: List of oligonucleotides used to clone the *NORAD* library.

Supplementary Tables ST9: Data relevant to Figure 6.

**Figure S1.**
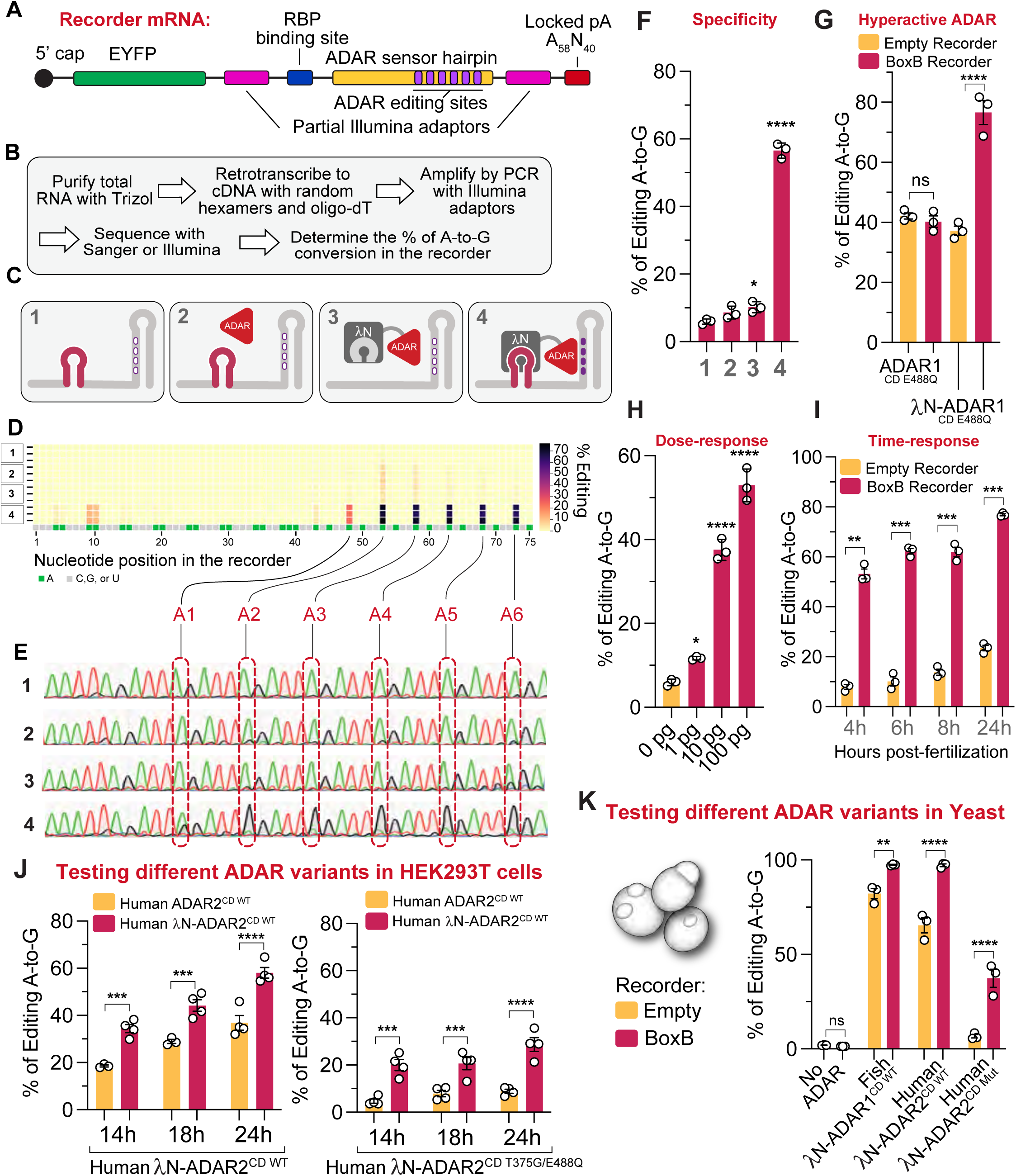
**A)** Cartoon depicting the organization of the recorder mRNA. **B)** Flowchart of the protocol RBPscan protocol from RNA extraction to determine the edits of a recorder. **C)** Cartoon describing the different combinations of writers and recorders to establish robust controls to test the specificity of RBPscan in the experiments conducted in zebrafish embryos upon injection of mRNAs. Similar to Figure 1C. **D)** Heat map showing the positions of nucleotides in the recorder hairpin, colored according to the percentage of editing. The wild-type catalytic domain of zebrafish ADAR1 was used. Recorder edits were inferred by Illumina sequencing, with the bar plot representation of this data shown in Figure 1D. Editing occurs preferentially at adenosines in mismatched positions. Samples are described in Figure S1C. **E)** Same samples as Figure1 D and S1D, but recorder edits are inferred from Sanger sequencing chromatograms. **F)** Bar plot indicating the % of editing triggered by ADAR^CD^ or λN22-ADAR^CD^ fusion in the corresponding recorders. Wild-type catalytic domain of zebrafish ADAR1 was used. Recorder edits are inferred by Sanger sequencing of the same samples of Figure 1D and described in Figure S1C. Error bars indicate □standard error of the mean of three technical replicates. White circles indicate the mean of individual replicates processed from pools of 30 zebrafish embryos each. *P* value from one-way ANOVA test with Dunnet’s multiple comparison test correction, \**p*=0.0374, \*\*\*\**p*<0.0001. **G)** Bar plot indicating the % of editing triggered by ADAR^CD^ ^E488Q^ or λN22-ADAR^CD^ ^E488Q^ fusion in the corresponding recorders. ADAR^CD^ ^E488Q^ is a hyperactive mutant of zebrafish ADAR1. Recorder edits are inferred by Sanger sequencing. Error bars indicate □standard error of the mean of three technical replicates. White circles indicate the mean of individual replicates processed from pools of 30 zebrafish embryos each. *P* value from one-way ANOVA test with Dunnet’s multiple comparison test correction, \*\*\*\**p*<0.0001. **H)** Bar plot indicating the % of editing triggered by increasing doses of λN22-ADAR^CD^ fusion in the BoxB recorder. Wild-type catalytic domain of zebrafish ADAR1 was used. Recorder edits are inferred by Sanger sequencing of the same samples of Figure 1F. Editing values for the 0 pg sample are reused from sample #1 (BoxB recorder only) of Figure S1F. Error bars indicate □standard error of the mean of three technical replicates. White circles indicate the mean of individual replicates processed from pools of 30 zebrafish embryos each. *P* value from one-way ANOVA test with Dunnet’s multiple comparison test correction, \**p*=0.0416, \*\*\*\**p*<0.0001. **I)** Bar plot indicating the % of editing triggered by λN22-ADAR^CD^ fusion in the empty or BoxB recorder at different time points after embryo fertilization. Wild-type catalytic domain of zebrafish ADAR1 was used. Recorder edits are inferred by Sanger sequencing of the same samples of Figure 1H. Error bars indicate □standard error of the mean of three technical replicates. White circles indicate the mean of individual replicates. *P* value from two-way ANOVA test with Sidak’s multiple comparison test correction, \*\**p*=0.0012, \*\*\**p*<0.001. **J)** Bar plot indicating the % of editing triggered by λN22 fusions with either ADAR^CD^ ^WT^ or ADAR^CD^ ^MUT^ in the BoxB recorder in HEK293T cells at different time points. In this experiment, ADAR^CD^ is derived from human ADAR2 wild-type or T375G/E488Q mutant. Recorder edits are inferred by Sanger sequencing. Error bars indicate □standard error of the mean of four technical replicates. White circles indicate the mean of individual replicates. Sample 24h λN22-ADAR2^CD^ ^T375G/E488Q^ is also shown in Figure 1I. *P* value from two-way ANOVA test with Sidaks’s multiple comparison test correction for ADAR^CD^ samples and mixed-effects analysis for ADAR^CD^ ^T375G/E488Q^, \*\*\**p*<0.001, \*\*\*\**p*<0.0001. **K)** Bar plot indicating the % of editing triggered by λN22-ADAR^CD^ of different origins in the empty or BoxB recorders in yeast. λN22-ADAR2^CD^ ^Mut^ stands for λN22-ADAR2^CD^ ^T375G/E488Q^. Recorder edits are inferred by Sanger. Error bars indicate □standard error of the mean of four technical replicates. White circles indicate the mean of individual replicates. λN22-ADAR2^CD^ ^T375G/E488Q^ are also shown in Figure 1L. *P* value from two-way ANOVA test with Sidaks’s multiple comparison test correction, \*\**p*<0.01, \*\*\*\**p*<0.0001.

**Figure S2.**
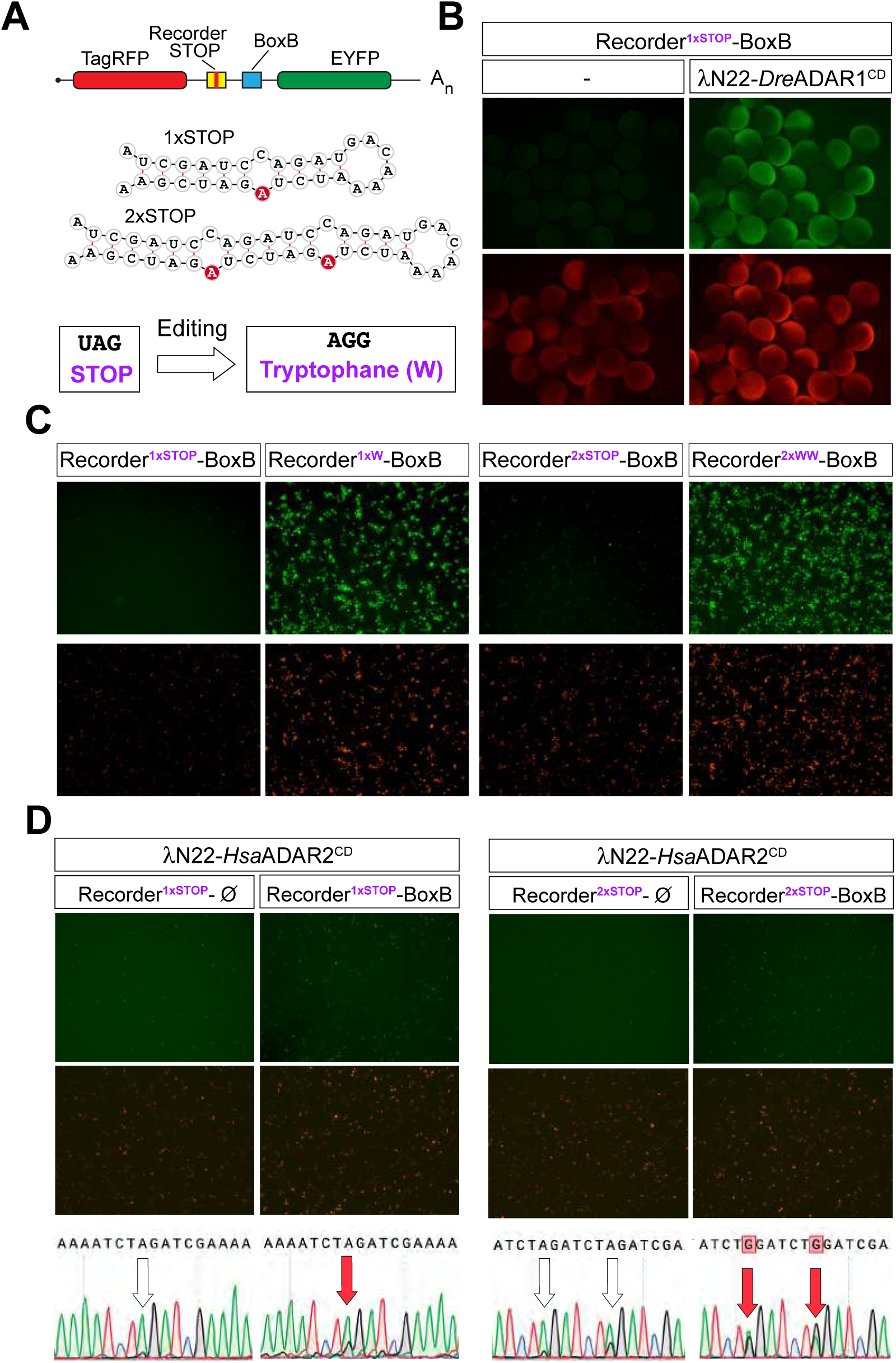
**A)** Cartoon illustrating the structure of the dual-fluorescence reporter, with the TagRFP and EYFP open reading frames separated by one or two stop codons in a hairpin and a BoxB site. **B)** Injection of the recorder in zebrafish embryos only produce green fluorescence if it is coinjected with λN22-*Dre*ADAR1^CD^. **C)** HEK293T cells are transfected with the recorder with one or two stops, or with the equivalent recorders with these stops sites fully edited to encode Tryptophane. **D)** HEK293T cells are transfected with the recorder with one stop next to a BoxB site or nothing, together with λN22-*Hsa*ADAR2^CD^. Sanger sequencing indicates the level of recorder editing in each condition. The experiment is repeated with the recorder containing two consecutive editable stop sites.

**Figure S3.**
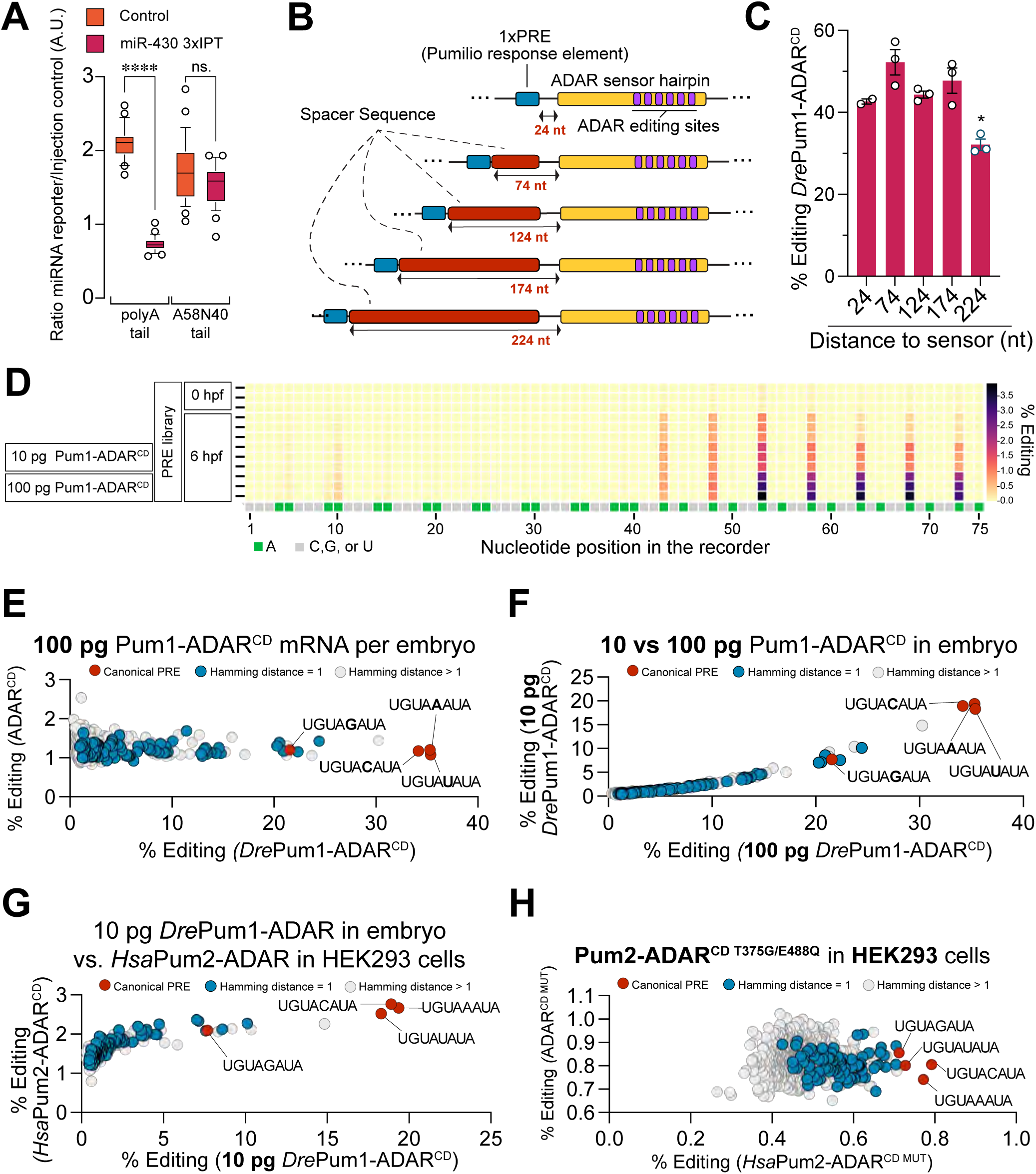
**A)** Box plots showing how the incorporation of a blocked polyA tail (58 A’s followed by 40 random nucleotides) to the recorder mRNA reduces its decay induced by miR-430 in the time frame tested (7 hours). Box plots enclose the 25th to 75th percentile and the whiskers delimit 5th to 95th percentiles, with the median indicated as a black line across the box. **B)** Cartoon illustrating a series of recorder constructs where the Pumilio response element (PRE) is pushed further apart from the recorder hairpin. The linker region was derived from zebrafish *Dicer1* 3’UTR deprived of PREs. **C)** Bar plot indicating the % of editing triggered by Pum1-ADAR^CD^ in the series or recorders with increasing spacer distance. Recorder edits are inferred by Sanger sequencing. Error bars indicate □standard error of the mean of three technical replicates. White circles indicate the mean of individual replicates processed from pools of 30 zebrafish embryos each. *P* value from one-way ANOVA test with Dunnet’s multiple comparison test correction, \**p*=0.0411. **D)** Heat map showing the positions and % of editing in PRE library induced by zebrafish Pum1-ADAR^CD^. Catalytic domain of wild-type zebrafish ADAR1 was used. **E)** Scatter plot showing the % of editing of individual motifs after co-injection into zebrafish embryos together with 100 pg of mRNA encoding ADAR^CD^ or zebrafish Pum1-ADAR^CD^. Catalytic domain of wild-type zebrafish ADAR1 was used. Each circle represents the mean editing of a given motif in three replicates. Circles are colored according to their Hamming distance to a canonical PRE. **F)** Scatter plot showing the correlation of motifs according the to the level of editing after injection in zebrafish with either 10 or 100 pg of mRNA encoding Pum1-ADAR^CD^. Source data from Figure 2K and S2E. **G)** Scatter plot showing the correlation of motifs according the to the level of editing Pum1-ADAR^CD^ (10 pg) in zebrafish embryos and Pum2-ADAR^CD^ in HEK293T cells. Source data from Figure 2K and 2L. **H)** Scatter plot showing the % of editing of individual motifs after transfection of HEK293T cells together with ADAR^CD^ ^T375G/E488Q^ or human Pum2-ADAR^CD^ ^T375G/E488Q^. Each circle represents the mean editing of a given motif in three replicates. Circles are colored according to their Hamming distance to a canonical PRE.

**Figure S4.**
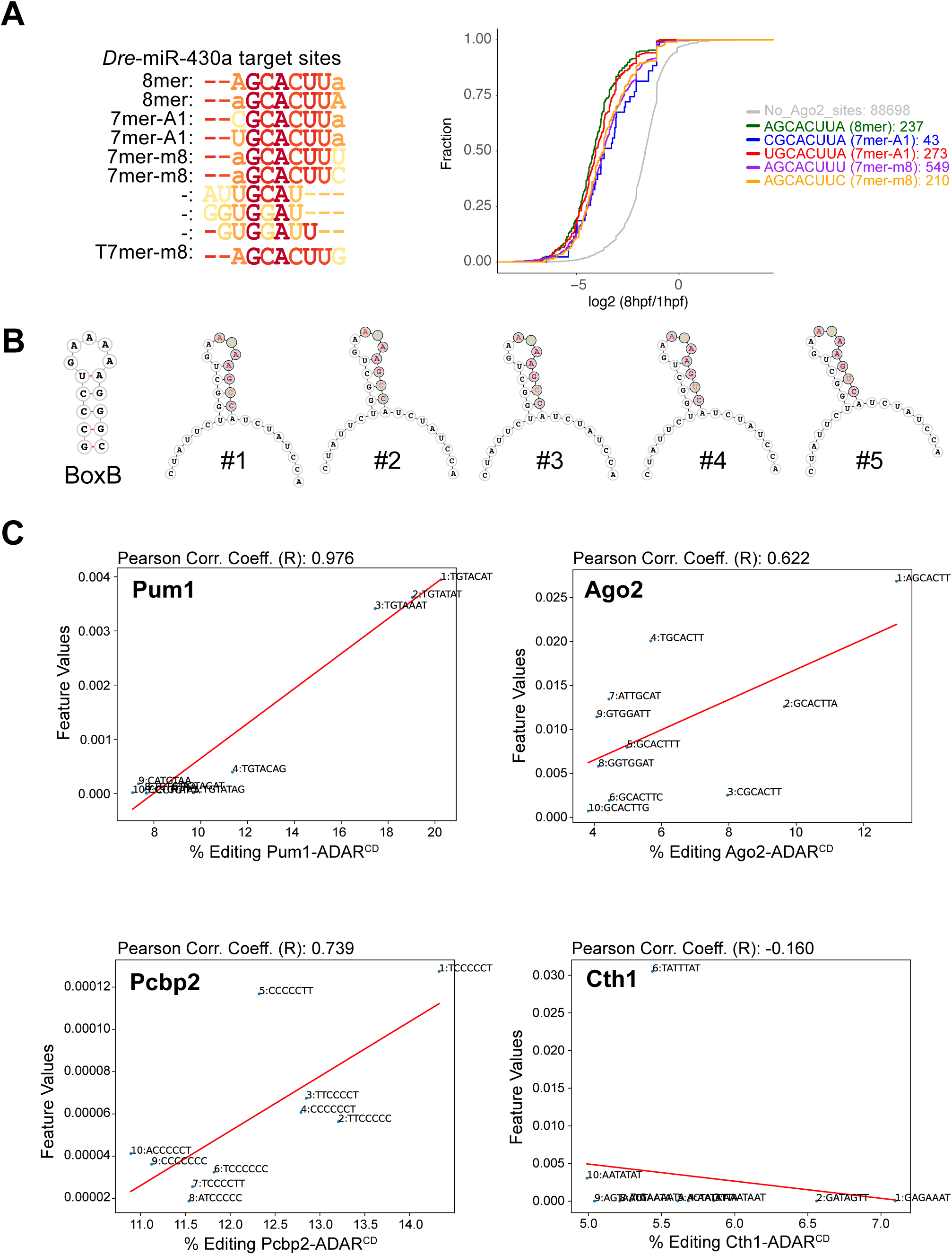
**A)** Sequence alignment showing the correspondence of the motifs enriched by RBPscan Ago2 and their complementarity to miR-430 seeds. Nucleotides are colored according to their prevalence in the alignment position. Lower cases indicate the nucleotides present in the fixed positions adjacent to the cloned motif. The alignment is also shown in Figure 4. Next, cumulative plots showing the degree of mRNA decay triggered by each miR-430 target site in a MPRA from Rabani *et al*^32^. **B)** RNA secondary structures of BoxB and the motifs enriched in the RBPscan screening with λN22-ADAR^CD^. In all instances, the structure is maintained independent of differences in the sequences. **C)** Correlation between the % of Editing measured in an RBPscan assay versus the calculated feature importance for decay predictions.

**Figure S5.**
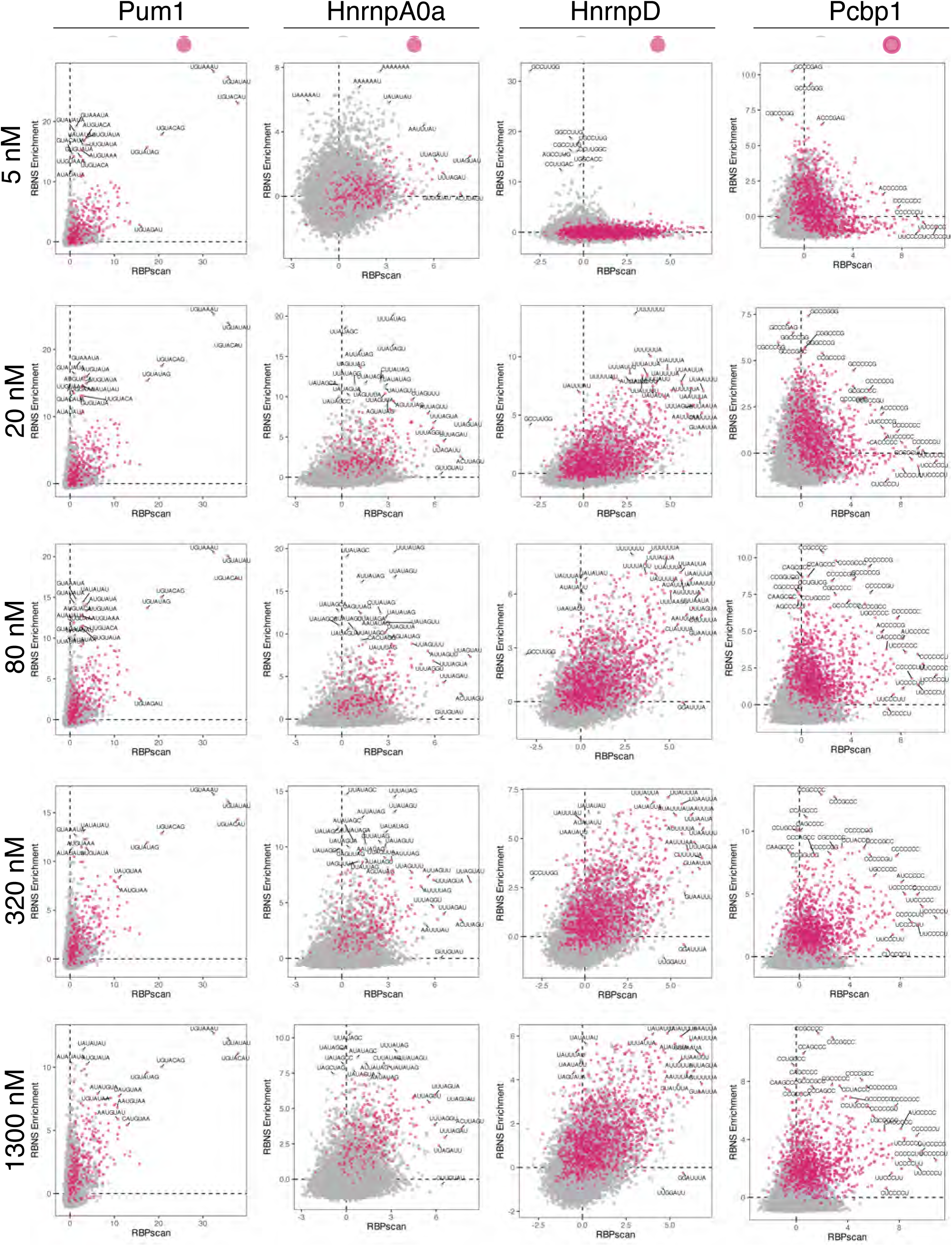
Benchmarking with Bind-n-Seq. **A)** Scatter plots showing the correlation between the enrichment of 7-mer motifs in Bind-n-Seq assay versus their editing efficiency in RBPscan. nM concentrations indicate the amount of purified truncated human protein used in the Bind-n-Seq assay. Red dots indicate the position of motifs containing the indicated motif. Bind-n-Seq data is extracted from Dominguez et al^15^.

**Figure S6.**
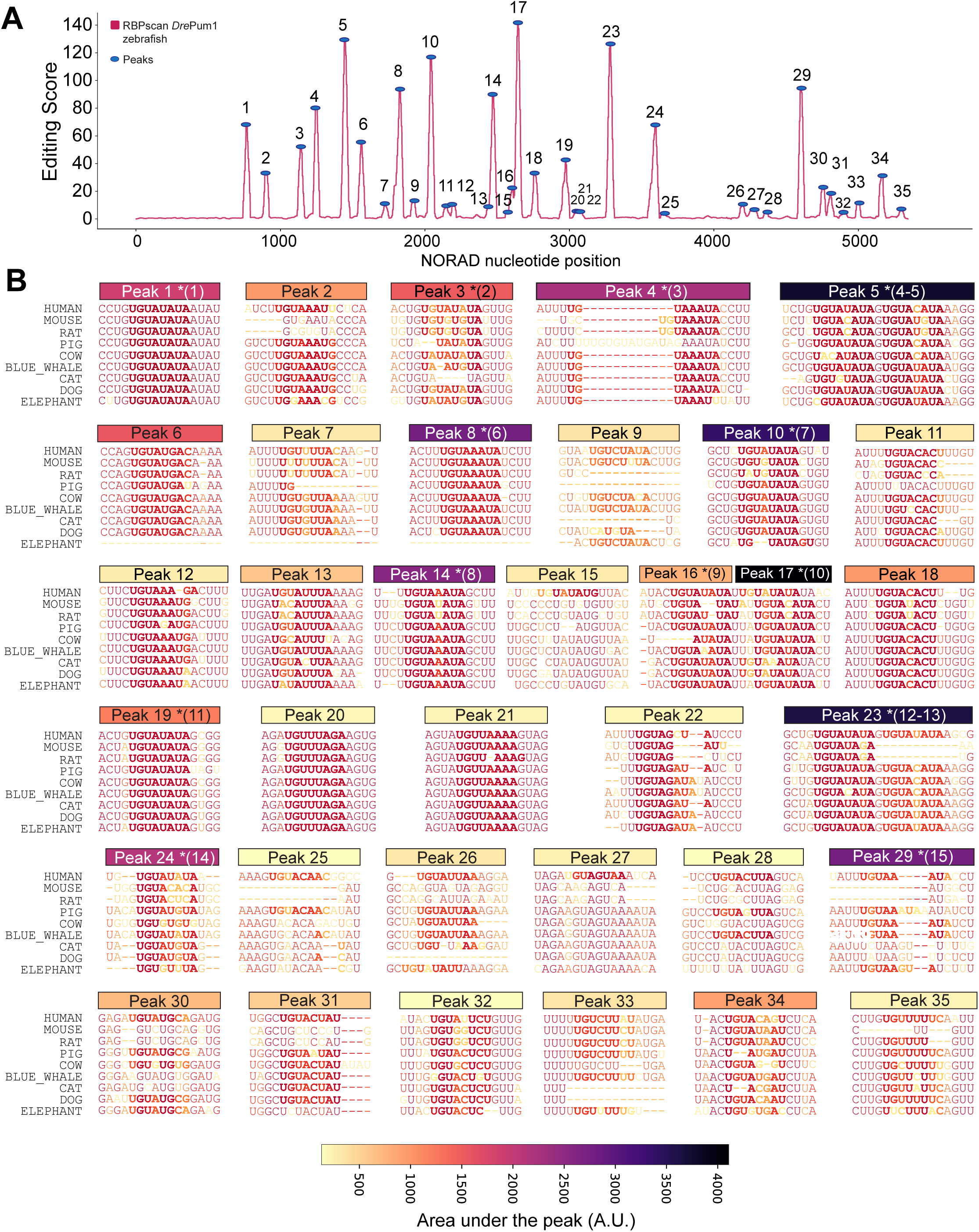
**A)** Editing score (red) after injection of the *NORAD* tilling recorder library in zebrafish embryos with *Dre*Pum1-ADAR^CD^. Same data as in Figure 5J, with additional blue circles indicate the peaks that have been annotated. **B)** *NORAD* alignments for the indicated species. The alignments depict Pum1-seed containing sequences that intersect with the annotated peaks Figure 5J. Peak number is inside a rectangle whose color reflects the value of the area under the corresponding peak. Asterisks indicate that the peak was previously described and the number in parenthesis indicates the peak number as annotated in Lee *et al*^40^.

## MATERIALS AND METHODS

### Plasmid generation

The plasmids encoding RBP-ADAR fusions were constructed by inserting the coding region of a full-length RBP tagged with a FLAG sequence at its N-terminus and the coding region of the catalytic domain of ADAR at its C-terminus into the pCS2+ plasmid at BamHI and XhoI using the NEBuilder HiFi DNA Assembly Kit (NEB #E2621L). The HA-tag sequence was used as a linker to separate the RBP and the catalytic domain of ADAR. For Ago2, the catalytic domain of ADAR was fused to its N-terminus. The coding regions of RBPs were PCR-amplified either from previously described plasmids^5^ or from cDNA synthesized from total RNA isolated either from 6-hour-old zebrafish embryos or from HEK293T cells. In all zebrafish experiments, the catalytic domain of wild-type ADAR1 from zebrafish (*Danio rerio*) was used to tag RBPs. For experiments in mammalian cells, RBPs were tagged with either the wild-type catalytic domain of human ADAR2 or the T375G/E488Q mutant, which was generated using Addgene plasmid #103870 as the PCR template. In yeast experiments, the λN22 peptide fused to the catalytic domain of wild-type zebrafish ADAR1, wild-type human ADAR2, and the T375G/E488Q mutant of human ADAR2 were compared. The plasmid encoding recorder mRNA was constructed by inserting the coding region of EYFP along with a 3′ UTR region containing a landing site with an EcoRI restriction site and a recorder hairpin structure (5’-TCCAATCCAATCCAATCCAATCCAATCCAATCCAATTAAATTAGATTAGATTAGATTAGATTAGATTAGATTAGA-3’) flanked by partial Illumina adapters (5’-ACACTCTTTCCCTACACGACGCTCTTCCGATCT-3’ and 5’-GTGACTGGAGTTCAGACGTGTGCTCTTCCGATCT-3’) into the pCS2+ plasmid using the NEBuilder HiFi DNA Assembly Kit (NEB). Additionally, a 58-nt poly(A) tail, PCR-amplified from Addgene plasmid #50562, was inserted into the recorder plasmid downstream of the recorder hairpin. All plasmids containing the genetically encoded poly(A) tail were transformed into and grown in the Stable competent *E. coli* strain (NEB #C3040H). To generate plasmids with specific binding motifs, the Recorder plasmid was linearized at the landing site using restriction enzyme EcoRI, and DNA oligos containing the motifs of interest were inserted using the NEBuilder HiFi DNA Assembly Kit (NEB).

### Generation of Pumilio recorder plasmid library

To prepare the input Recorder plasmid library for the Pumilio experiment, a pool of 1000 oligonucleotides (PRE_library_oligos) was ordered from Twist Bioscience. Each oligonucleotide contains a unique 8-mer sequence and flanking sequences necessary for NEBuilder HiFi DNA Assembly into the Recorder plasmid. The Pumilio oligo library was designed to include 4 perfect Pumilio Response Elements (PREs) (TGTANATA), 88 variants differing by 1 nucleotide from the perfect PREs (Hamming distance = 1), 844 sequences differing by 2 nucleotides from the perfect PREs (Hamming distance = 2), 40 sequences with a Hamming distance of 8 from the perfect PREs, and 24 manually selected sequences representing homopolymers and partial PREs (see Table_S3 for all sequences). To generate a library of recorder plasmids with different sequence variants, NEBuilder High-Fidelity Assembly was performed using a recorder plasmid linearized by EcoRI and a PRE_library_oligos. The resulting ligation mixture was transformed into Stable *E. coli* and plated on ten 150 mm agar plates containing ampicillin (100 μg/μl). More than 30,000 individual colonies were collected from several plates using a cell scraper. The bacteria were pelleted by centrifugation, and plasmid DNA was purified using the PureLink HiPure Plasmid Miniprep Kit (Thermo Fisher #K210002). Equal representation of all sequence variants was confirmed by next-generation sequencing.

### Generation of 7N recorder plasmid library

To prepare the input 7N plasmid library for the motif discovery experiment, an oligonucleotide pool (7N_library_oligos) containing random nucleotides at 7 positions and flanking sequences for NEBuilder HiFi DNA Assembly was ordered from IDT (a total of 16,384 sequences). A constant adenosine was incorporated directly after the 7^th^ random nucleotide to enable the identification of complete perfect PREs (8-nt long) or 8-mer binding motifs of microRNAs, which preferentially bind to sites that have adenosine at the first position. To generate a library of recorder plasmids with different sequence variants, NEBuilder High-Fidelity Assembly was performed using a recorder plasmid linearized by EcoRI and a 7N_library_oligos. The resulting ligation mixture was transformed into Stable *E. coli* and plated on twenty-five 150 mm agar plates containing ampicillin (100 μg/μl). More than 70,000 individual colonies were collected from several plates using a cell scraper. The bacteria were pelleted by centrifugation, and plasmid DNA was purified using the PureLink HiPure Plasmid Miniprep Kit (Thermo Fisher #K210002). Equal representation of all sequence variants was confirmed by next-generation sequencing.

### Generation of NORAD recorder plasmid library

To prepare the input Recorder plasmid library for the NORAD experiment, a pool of tiled oligonucleotides spanning the sequences of *hsNORAD* RNA (see Table_S8 for all sequences) was ordered from Twist Bioscience. To construct the oligonucleotide library covering the entire sequence of *hsNORAD* RNA, we designed a pool of overlapping oligonucleotides. Each oligonucleotide was 30 nt in length, with a 20-nt overlap between consecutive sequences creating a continuous coverage of the entire *hsNORAD*. To generate a library of recorder plasmids with *NORAD*-spanning sequences, NEBuilder High-Fidelity Assembly was performed using a recorder plasmid linearized by EcoRI and a NORAD_library_oligos. The rest of library cloning procedure was performed as described for Pumilio recorder plasmid library.

### In vitro transcription

The recorder- and writer-encoding plasmids and libraries (∼3 µg) were linearized using the NotI restriction enzyme in 1X CutSmart Buffer for 2 hours at 37 °C. All Writer-encoding plasmids retain an SV40 polyadenylation site after linearization, while Recorder plasmids with a genetically encoded poly(A) tail have an additional 40 nucleotides at the 3’ end to reduce mRNA deadenylation. The linearized DNA fragments were purified using the Monarch PCR & DNA Cleanup Kit (NEB #T1130L). Next, 1 µg of linearized DNA was used for an in vitro transcription reaction performed with the mMESSAGE mMACHINE SP6 Transcription Kit (ThermoFisher #AM1340) at 37 °C for 2 hours. The resulting RNA was purified using the RNA Clean & Concentrator-5 Kit (Zymo Research #R1014).

### Zebrafish strains

Zebrafish strains were bred, handled, and maintained according to the standard laboratory conditions under IACUC protocol PROTO201800373 at Boston University. Experiments were performed in hybrid wild-type strain crosses obtained from AB/TU and TL/NIHGRI breeders. The sex of zebrafish embryos was not considered because sex determination occurs at later time points of development.

### Microinjections into zebrafish embryos

Microinjections into embryos were performed as described previously^44^. Briefly, one-cell stage zebrafish embryos were collected and microinjected with glass needle into the animal pole. Needles were calibrated to inject 1 nL. Immediately after injection, embryos were transferred into an agarose-coated 100 mm Petri dish and incubated at 28.5° C until processing. 30-40 embryos were collected per replicate at the indicated time points.

### Cell culture and Transfection into HEK cells

HEK293T cell lines were maintained in humidified 5% CO_2_ incubator at 37°C in DMEM high glucose media (4.5 g/L) (Corning #MT10013CV) containing 10% FBS, 1% sodium pyruvate, penicillin/streptomycin, 2 mM L-glutamine. For routine passaging cells were washed with 1X DPBS (Phosphate buffered saline, ThermoFisher # 14190144) and dissociated using 0.25% trypsin-EDTA (ThermoFisher # 25200056). For the experiments described in Figure 1, 1 μg of writer and recorder plasmids were transfected into cells seeded in 12-well plates at 70% confluency using Lipofectamine 3000 Reagent (ThermoFisher #L3000008). The cells were collected at the indicated time points post-transfection and were washed twice with 1X DPBS before freezing. For the experiments described in Figures 2 and 5, 2.5 μg per well of writer plasmids were first transfected into 6-well plates at 50% confluency using Lipofectamine 3000 reagent. Twelve hours later, 3 μg per well of the recorder mRNA library was transfected into the same cells using MessengerMAX transfection reagent (ThermoFisher #LMRNA003). Cells were collected 24 hours after RNA transfection, washed twice with 1X DPBS, and then frozen.

### RBPscan in yeast

The fusions of λN22 with zebrafish ADAR1, human ADAR2, and human T375G/E488Q ADAR2 were cloned into the pATP425 yeast (*Saccharomyces cerevisiae*) expression vector using the NEBuilder HiFi DNA Assembly Cloning Kit. A single colony of *S. cerevisiae* was cultured overnight at 28 °C and 200 rpm in 5 ml yeast extract-peptone-dextrose (YPD) medium (TaKaRa #630409). The overnight culture was then diluted to an optical density (OD600) of 0.1–0.2 in a total volume of 10 ml of YPD. Following dilution, the cells were grown at 28 °C and 200 rpm until the OD600 reached 0.6. The cells were centrifuged at 2000 rpm for 5 min, washed twice with water, and then resuspended in 100 µl of 1X Tris-EDTA and 1X lithium acetate (TE/LiAc, pH 7.5). The plasmid encoding the writer and recorder was transformed using the standard LiAc/ss-DNA/PEG protocol. The transformation reactions were plated on Minimal SD Base (TaKaRa #630411) leucine drop-out plates and incubated at 28 °C for 48 hours. EYFP-positive colonies were selected and inoculated into 5 ml of SD Base (-Leu) medium, then grown overnight at 28 °C and 200 rpm. EYFP-positive colonies were selected and inoculated into 5 ml of SD Base (-Leu) medium, then grown overnight at 28 °C and 200 rpm. The overnight culture was diluted to an optical density (OD600) of 0.1–0.2 in a total volume of 10 ml of SD Base (-Leu) and grown until the OD600 reached 0.6. The cells were pelleted by centrifugation at 3000 rpm for 5 minutes, resuspended in 1 ml of water, and transferred to a 1.5 ml tube. The cells were washed twice with water and frozen. Cell pellets were lysed in 400 µl of yeast lysis buffer (LETS buffer: 0.1 M LiCl, 0.01 M EDTA, 0.01 M Tris-HCl, pH 7.4, 0.2% SDS) mixed with 400 µl of acid phenol:chloroform (pH 4.5, ThermoFisher #AM9722) and filled with glass beads (with beads amount equal to one full PCR tube) (Sigma #G8772-100G). The mixture was vortexed six times for 30 seconds each, with samples kept on ice between each round of vortexing. A total of 250 µl of the supernatant was collected for total RNA purification by centrifugation at 13,000 rpm for 10 minutes at 4 °C. The collected supernatant was processed directly using the Direct-zol RNA Miniprep Kit (Zymo Research #R2050), following the instructions.

### RNA extraction and Reverse Transcription (RT) reaction

Total RNA from zebrafish or cells was extracted using 1 mL of TRIzol (ThermoFisher #15596026) per sample. RNA from the aqueous phase was precipitated with one volume of isopropanol and subsequently washed once with 80% ethanol. RNA pellet was resuspended in 20 □L of water. Reverse transcription was performed with 1 μg of total RNA in 20 □L of total volume using the LunaScript RT SuperMix Kit (NEB #E3010) under the following conditions: 2 minutes at 25°C, 10 minutes at 55°C, and 1 minute at 95°C.

### Analysis of RNA editing using Sanger sequencing

To analyze the A-to-G conversion in the recorder hairpin, a PCR reaction was performed to amplify the 3’UTR region of the recorder mRNA, which contains the tested binding sequence and the recorder hairpin. Briefly, 1 µL of cDNA was used for the PCR reaction with rec_forward and rec_reverse primers. For SV40-containing plasmids, the rec_reverse_SV40 primer was used, and for plasmids with a genetically encoded poly(A) tail, the rec_reverse_A58N40 was used (see Table_S1). The reaction was performed using Q5 High-Fidelity MasterMix (NEB # M0492) in a total volume of 25 µL for 35 cycles at 98 °C for 15 seconds, 63 °C for 20 seconds, and 72 °C for 30 seconds. PCR products were resolved on a 2% agarose gel in 1X TAE buffer, and DNA bands corresponding to the expected product size were purified using the Monarch DNA Gel Extraction Kit (NEB #T1120). The purified DNA products were then analyzed by Sanger sequencing using the rec_forward PCR primer. The intensity of chromatographic peaks for adenosine and guanosine at specific positions was quantified using the EditR tool^45^.

### Library preparation and next-generation sequencing

To prepare a library of for next-generation sequence analysis, a PCR reaction was performed to amplify the 3’UTR region of the recorder mRNA, which contains the tested binding sequence and the recorder hairpin. Partial Illumina adapters present in the recorder mRNA are complementary to parts of the i5 and i7 TruSeq Illumina primers, which also contain unique barcodes and sequencing adapters (see Table_S1 to see all sequences). Briefly, 2 µL of cDNA was used for the PCR reaction with i5 and i7 TruSeq Illumina primers (see Table S1 for the complete list of primers). The reaction was performed using Q5 High-Fidelity MasterMix (NEB # M0492) in a total volume of 25 µL for 12-18 cycles at the following settings 98 °C for 15 seconds, 63 °C for 20 seconds, and 72 °C for 30 seconds. The number of cycles was optimized to provide enough material for sequencing while minimizing potential bias due to PCR overamplification. PCR products were resolved on a 2% agarose gel in 1X TAE buffer, and DNA bands corresponding to the expected product size were purified using the Monarch DNA Gel Extraction Kit (NEB #T1120). Concertation of isolated DNA products was quantified using Qubit dsDNA Quantification Assay Kit (ThermoFisher #Q32853). Several libraries were pooled together at the equimolar concertation. Next-generation sequencing was performed on the NovaSeq X Plus Illumina platform using a 150-base pair end cycle at 10-20 million reads per library.

### Analysis of RBPscan data

First, paired-end reads were merged using the BBMap suite, and then all FASTQ files were converted into TXT files. RBPscan data were analyzed using a custom Python script. First, only reads containing intact sequences located between Illumina adapters were filtered. G substitutions at adenosine sites in the recorder hairpin were allowed to account for both edited (“G”) and non-edited (“A”) positions in each read. Then, all variable sequence motifs were identified and counted. The threshold of 1000 reads per motifs was used for PREs and NORAD libraries and 10 reads for 7N library. Then, we counted all A-to-G substitutions for each motif in each read at each adenosine position to estimate the editing efficiency induced by the presence of the motifs. Then mean percent of A-to-G editing was calculated for all reads with same motif and editing efficacy score.

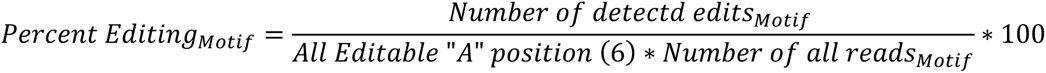

Each experiment was performed in three replicates and the mean percent of editing was represented on the graphs.

### Analysis of NORAD libraries

The mean percent editing of NORAD-tiling sequences was calculated as described in the “Analysis of RBPscan Data” section. Subsequently, cumulative percent editing (RBPscan score) was determined for each nucleotide within the NORAD genomic sequence. This was achieved by summing up the percent editing values for each nucleotide of the NORAD sequences across all reads that include the respective nucleotide, using a custom Python script. The sequencing coverage of NORAD genomic sequences was expressed as reads per million (RPM) values. For normalization, the RBPscan scores of either the ADARcd condition or the no-protein control were subtracted from the RBPscan scores obtained for Pumilio (PUM1 or PUM2).

### Analytic Pearson residuals analysis

For Ago2-Adar miR-430 canonical target experiments, total edits per motif were normalized by sequencing depth following the method defined previously^46^. First, the expected counts were estimated as the product of the occurrence of each recorder and a rate parameter. The rate parameter was calculated as the mean of total edits divided by the occurrences across all recorders. Pearson residuals were then calculated using a negative binomial model, with the dispersion parameter set to 5. This approach normalizes the observed edit counts by their expected values, accounting for abundance due to variability in sequencing depth, microRNA-mediated repression, and overdispersion in the data.

### Calculation of 7-mer enrichment values from RNA Bind-n-seq

For each protein concentration and its respective input library, analysis is performed following the ENCODE RNA Bind-n-Seq computational pipeline and the methods previously described^15,47^. A custom script for analysis was written in Python and C for calculation 7-mer enrichment values. Briefly, the total occurrences of each 7-mer across all reads in the library were counted. These counts were normalized to frequencies by dividing each 7-mer count by the total number of possible 7-mers, calculated as (sequence length−k+1)*X, where k=7 and X is the total number of reads in the library. This normalization was performed independently for the protein-bound libraries and the input libraries. The enrichment for each -7mer was then computed as the ratio of its frequency in the protein library to its frequency in the input library. The enrichment values for each 7-mer were compared to corresponding editing values obtained from RBPscan. For each Bind-n-seq protein concentration the enrichment values are compared to RBPscan’s percent editing by calculating the spearman correlation.

### Analysis of MPRA data

The RNA stability data used in Figures 4G and H, S3C, and S4D were derived from Rabani *et al*, 2017^32^. PREs were identified and counted in all sequences that were analyzed in the library. In cumulative plots represented in Figure 4G and H the control group contains all PRE elements that showed strong binding to PUM1 (UGUAAAUA, UGUAUAUA, UGUACAUA, UGUAGAUA, UGUACAGU, UGUACAGA, UGUACAGG, UGUAUAGU, UGUAAAUU, UGUAUAGA). In Figure S3C control group excluded all analyzed miR-430 binding sites (AGCACUUA, CGCACUUA, UGCACUUA, AGCACUUU, AGCACUUC). Cumulative plots were generated using the R package stat_ecdf.

### Calculation of RBP motif importance in MPRA data using SHapley Additive exPlanations (SHAP value)

To measure the correlation between the mean A-to-G percent editing for each RBP and its effect on RNA stability obtained from MPRA performed in zebrafish embryos^32^, we try to assign each RBP motif a feature importance based on a machine learning algorithm. We hypothesize that motifs with high percent editing are high indicators of RNA stability, thus having a motif with a high percent editing will have a higher effect on the RNA stability. To do this, we train an XGBoost Regressor, with RBP motif sequence composition as input and RNA stability as output. After we obtain a model capable of predicting RNA stability with Pearson correlation coefficient ranging from 0.52 to 0.62, we utilize SHAP values to extract the importance of each feature, in this case, a motif. We then plot this newly obtained feature importance against mean A-to-G percent editing and measure their Pearson Correlation Coefficient. A strong correlation supports our hypothesis by showing that motifs with high percent editing show large indication of RNA stability.

### Detection of RBP binding using fluorescent readout

For the experiments described in Figure S2, 1 μg of writer and fluorescent recorder plasmids were transfected into HEK293T cells seeded in 12-well plates at 70% confluency using Lipofectamine 3000 Reagent (ThermoFisher #L3000008). The imaging of life cells was performed 24 hours after transfection using Discovery V12 Fluorescence Stereomicroscope (Zeiss).

### Analysis of eCLIP PUM1 and PUM2 data

The eCLIP data for hsaPUM1 and hsaPUM2 (ENCFF231WHF.bam and ENCFF732EQX.bam performed in K562 cell were obtained from ENCODE eCLIP dataset^47^ (Accession ID: ENCFF064COB.bam, ENCFF583QFB.bam ENCFF231WHF.bam, ENCFF732EQX.bam respectively). Read coverage over the genomic region was extracted using a custom Python script.

### Statistical analysis

All statistical analysis documented here were conducted with GraphPad Prism, v.10.0.3. Figure legends describe the sample size and statistical analysis done in each corresponding plot. As a general approach, to determine the statistical significance of differences between two groups we conducted two-tailed unpaired t-test while For multiple groups we conducted a one-way ANOVA test followed with a Dunnett’s multiple comparisons test. In all instances, the significance threshold was set for *p* < 0.05. R package “drc” was used to plot Pum1 dose-response curves and apparent relative *K_D_*’s.

### Use of generative AI-assisted technologies

During the preparation of this work, DC used ChatGPT 4o in order to improve the grammar and style of the manuscript. After using this tool/service, the DC and DK reviewed and edited the content as needed and take full responsibility for the content of the publication.

